# Integrative Transcriptomics Reveals Layer 1 Astrocytes Altered in Schizophrenia

**DOI:** 10.1101/2024.06.27.601103

**Authors:** Julio Leon, Satoshi Yoshinaga, Mizuki Hino, Atsuko Nagaoka, Yoshinari Ando, Jonathan Moody, Miki Kojima, Ayako Kitazawa, Kanehiro Hayashi, Kazunori Nakajima, Carlo Condello, Piero Carninci, Yasuto Kunii, Chung Chau Hon, Jay W. Shin, Ken-ichiro Kubo

## Abstract

Schizophrenia is one of the most prevalent psychiatric disorders with unclear pathophysiology despite a century-long history of intense research. Schizophrenia affects multiple networks across different brain regions. The anterior cingulate cortex (ACC) is the region that connects the limbic system to cognitive areas such as the prefrontal cortex and represents a pivotal region for the etiology of schizophrenia; however, the molecular pathology, considering its cellular and anatomical complexity, is not well understood. Here, we performed an integrative analysis of spatial and single-nucleus transcriptomics of the postmortem ACC of people with schizophrenia, together with a thorough histological analysis. The data revealed major transcriptomics signatures altered in schizophrenia, pointing at the dysregulation of glial cells, primarily in astrocytes. We further discovered a decrease in the cellular density and abundance of processes of interlaminar astrocytes, a subpopulation of astrocytes specific to primates that localize in the layer 1 and influence the superficial cortical microenvironment across layer 1 and layers 2/3 of the cortex. Our study suggests that aberrant changes in interlaminar astrocytes could explain the cell-to-cell circuit alterations found in schizophrenia and represent novel therapeutic targets to ameliorate schizophrenia-associated dysfunction.

## Main

Schizophrenia (SCZ) is a complex psychiatric disorder that affects about 1% of people worldwide. It presents a substantial challenge to mental health systems, primarily due to its multifaceted nature and the often-limited effectiveness of current antipsychotic treatments^1^. The quest to unravel the pathophysiological foundations of SCZ has historically been arduous because there are no specific hallmarks in standard histology, unlike neurodegenerative diseases such as Alzheimer’s disease, leading to its description as the ‘graveyard of neuropathologists’^2^. We postulate that the pathophysiology can be found at the molecular and cellular levels if we adequately address the complexity and heterogeneity of the human brain, particularly in terms of cell composition and spatial wiring.

Single-cell technologies are helping to address this heterogeneity. Previous studies have identified specific transcriptomic alterations associated with neural cell subtypes in the prefrontal cortex of SCZ patients^3–5^. However, little is known about the key molecular changes in the microenvironment within the layers of the cortex affecting not only neurons but also glial cells forming interconnected networks in critical brain regions affected in SCZ. The anterior cingulate cortex (ACC) is crucial for the pathophysiology of SCZ. It connects the phylogenetically-old ‘emotional’ limbic system, such as the hippocampus, to the evolutionary-expanded ‘cognitive’ prefrontal cortex^6^, and plays an important role in the way we evaluate the reality of our environment (prediction error estimation)^7,8^. The dysfunction of this region of the brain is associated with SCZ symptomatology^9,10^. Recent brain imaging studies of SCZ have identified altered brain networks in the ACC^11^, including reduced gray matter volume that uniquely appears both prior to and following the onset of full-blown psychosis, encompassing stages from clinical high risk to chronic schizophrenia^12^. Microscopically, these structural changes coincide with an alteration in the density of von Economo neurons^13^ and interstitial white matter neurons^14^. Furthermore, while most previous research has predominantly concentrated on neurons, studies investigating glial cells within the context of schizophrenia in the ACC have yielded inconclusive results^15^. These inconsistencies arise, in part, because these studies do not consider the cellular diversity and anatomical complexity of the ACC.

To bridge this gap, our study employs an unprecedented integration of Visium spatial transcriptomics (ST), single-nucleus RNA sequencing (sn-RNA-seq), RNA fluorescence *in situ* hybridization (FISH), and immunohistochemistry (IHC) to reveal the molecular landscape of the ACC in schizophrenia (SCZ). To our knowledge, this is the first study to characterize and analyze the complexities of this important region, the ACC, in SCZ at single-cell resolution while considering its spatial context.

Spatial transcriptomics enabled us to identify, unbiasedly, cortical layers within the ACC and other domains associated with meninges or the blood vessels. Following the integration with sn-RNA-seq data and statistical comparisons by cortical layers, we observed a significant influence of glial cell signatures, particularly astrocytes, on the transcriptomic variations associated with SCZ. Further histological investigations revealed that a primate-specific type of astrocytes, interlaminar astrocytes (ILAs) that originate in layer 1 and extend long processes deep into cortical layers, exhibited reduced cell density and abundance of the processes in SCZ. Notably, we discovered that these alterations were influenced by sex and correlated with disease duration. In addition, we found a negative relationship between ILA processes and GFAP+ cell density in the grey matter in aging. Our findings suggest the critical role of astrocytes in the disrupted brain circuitry of SCZ and propose a novel morphological marker for its histopathological diagnosis.

## Results

### Spatial transcriptomics reveals molecular cortical domains from the ACC

To capture molecular alterations in the ACC of SCZ at single-cell and spatial levels, we captured nuclear and spatial gene expression features from the same fresh-frozen ACC tissue blocks, which included the gray matter (GM) and white matter (WM). ACC samples included six control donors without history of psychiatric or neurological disorders and six samples from patients clinically diagnosed with SCZ (Extended Data Table 1). We used the same fresh-frozen and additional formalin-fixed paraffin-embedded (FFPE) tissue blocks for RNA-FISH and IHC, respectively (Fig. 1a). Following data processing and batch correction, we analyzed 29,935 nuclei and identified eight major cell types in the ACC using the Azimuth package (Methods) (Fig. 1b). We confirmed this annotation by the enrichment of the major cell type marker genes, as indicated in Fig. 1c. As for Visium ST, we performed QC, data integration, and clustering and unveiled nine major clusters across all samples (Figs. 1d-f and Extended Data Figs. 1a-d), each exhibiting distinct gene expression patterns and anatomical localizations. Cluster 5, residing in cortical layer 1 (L1), displayed elevated levels of L1-specific genes *AQP4* and *AGT*. Cluster 3, found in the superficial cortical layers (L2/3), was enriched with *CCK* and *ENC1* genes. Clusters 0 and 6, corresponding to the deep cortical layers (L5/6) expressing *STMN2* and *DIRAS2*^16^, were identified. Clusters 1 and 2, primarily located in the WM, were marked by a high expression of the *MBP* gene. Cluster 7, situated at the GM and WM interface, was labeled as the ‘Border’ cluster, corresponding to the L6b. Cluster 8, associated with the choroid plexus and expressing genes such as *APOD*, *B2M*, *TAGLN*, and *IGFBP7*, was designated the Meninges Associated Cluster (MAC). Lastly, Cluster 4, enriched with hemoglobin genes *HBB*, *HBA2*, and *HBA1*, was identified as a Vascular Associated Cluster (VAC). Cortical layers and other domains showed strong concordance with the nuances of the human cortical anatomy by which some ST spots will contain gene signatures of the neighboring layers reflecting the intricate physical interaction of cell-to-cell networks^17^ (illustrated in Extended Data Fig. 1b). Our integration of sn-RNA-seq and ST data comprehensively delineates the cellular and histological complexity of human ACC tissues. This detailed molecular neuroanatomy served as a crucial foundation for subsequent analysis by cortical domain and cell type, setting the stage for detecting specific alterations associated with SCZ.

**Fig. 1.**
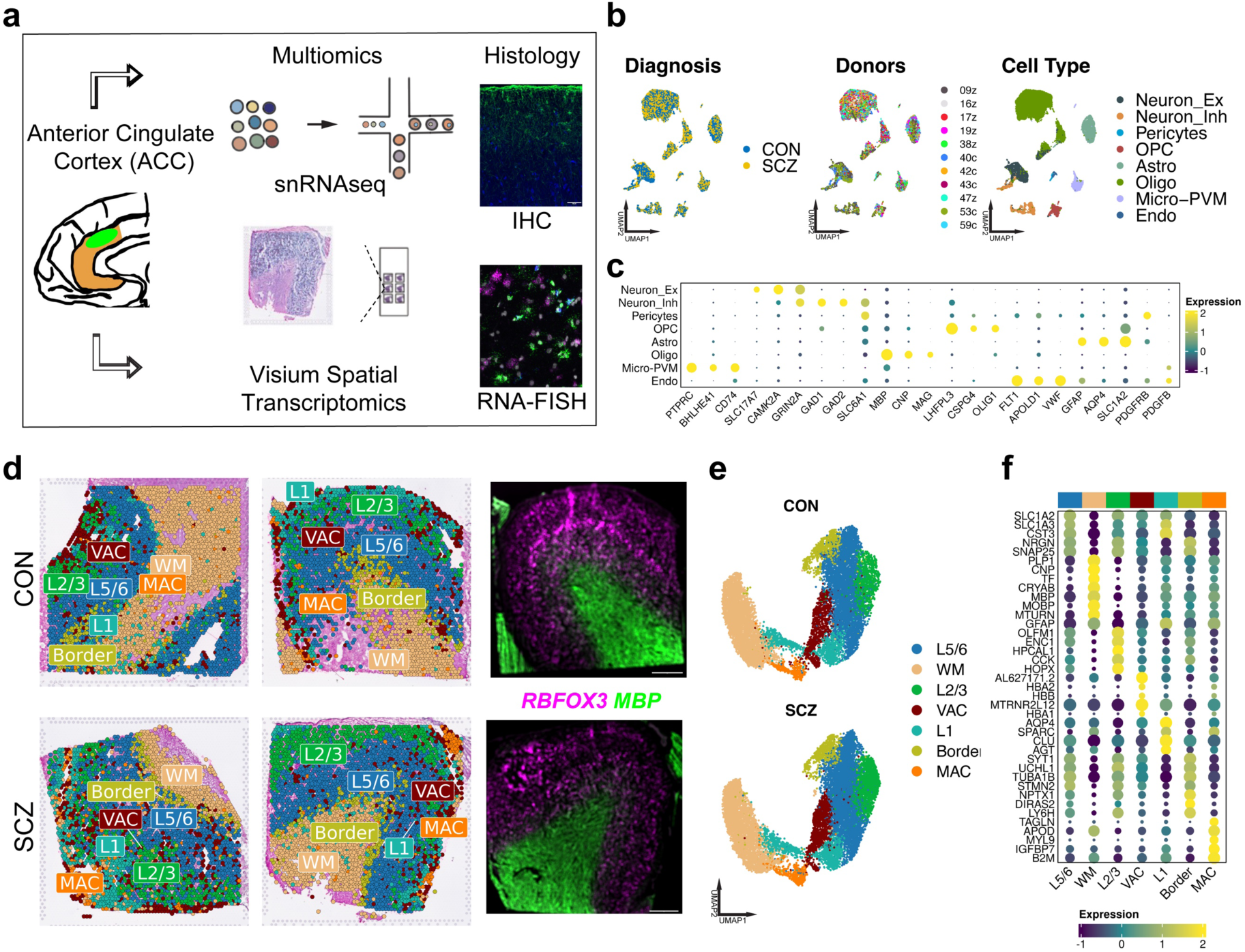
Single nuclei and spatial transcriptomics integrative analysis from the ACC of SCZ and control donors. **a**, Scheme for the integrative analysis of the ACC region in CON and SCZ. Samples were analyzed using sn-RNA-seq and spatial transcriptomics. Later, findings were confirmed in larger cohorts using RNA-FISH (5 CON and 5 SCZ samples) and IHC (13 CON and 17 SCZ samples). **b**, Post-integration uniform manifold approximation and projection (UMAP) plots of sn-RNA-seq profiles by diagnosis, donors and cell type. Neuron_Ex, excitatory neurons; Neuron_Inh, inhibitory interneurons; Pericytes; OPC, oligodendrocyte progenitor cells; Astro, astrocytes; Oligo, oligodendrocytes; Micro-PVM, microglia and perivascular macrophages. **c**, Cell-type markers confirming cell-type assignments. **d**, ACC domains identified by unbiased clustering and classified based on histological location and gene expression signatures, mapped onto H&E staining. RNA-FISH for *RBFOX3* (neurons) and *MBP* (oligodendrocytes) of quasi-adjacent sections is shown after averaging intensity in 55 μm to mimic Visium ST. Scale bars, 1 mm. L5/6, cortical layers 5 and 6; WM, the white matter; L2/3, cortical layers 2 and 3; VAC, vascular associated cluster; L1, cortical layer 1; Border; MAC, meninges associated cluster. **e**, UMAP plots of spatial transcriptomics cortical domains. **f**, ACC domain’s main gene expression signatures.

### Spatial transcriptomic and pathway signature changes in the ACC of patients with SCZ

To comprehensively identify brain-region-dependent dysregulated genes in the ACC of SCZ patients, we executed a differential gene expression (DGE) analysis. We used a logistic regression model and excluded mitochondrial and ribosomal RNA genes to minimize technical biases. Our analysis revealed 344 dysregulated genes across various cortical layers, with the largest number in the L2/3 exhibiting 99 dysregulated genes. This may suggest an enhanced vulnerability of L2/3 compared to other regions. On the other hand, the Border and WM regions displayed significant dysregulation of 73 and 56 genes, respectively (Fig. 2a, Table 1). A detailed analysis of genes showing significant dysregulation (adjusted *p* < 0.05, fold change > 1.3) (Fig. 2e) across cortical layers showed the downregulation of the metallothionein (*MT*) *MT1* and *MT2* gene family in L2/3 and L5/6, known for zinc ion binding and stress regulation that are primarily expressed in astrocytes^18^, and the *MBP* gene, critical for myelin formation^19^. Conversely, we found the upregulation of immune response genes including *CD74*, *HLA-B*, *HLA-DRA*, and the *IFI6* interferon response gene in the L1, L2/3, and L5/6. Remarkably, the *GFAP* gene, indicative of astrocytic activity^20^, was upregulated across all cortical regions, alongside *VIM* in all regions except in the VAC in SCZ compared to control. In addition, we detected the upregulation of *SERPINA3* in the L5/6, Border, and WM.

**Fig. 2.**
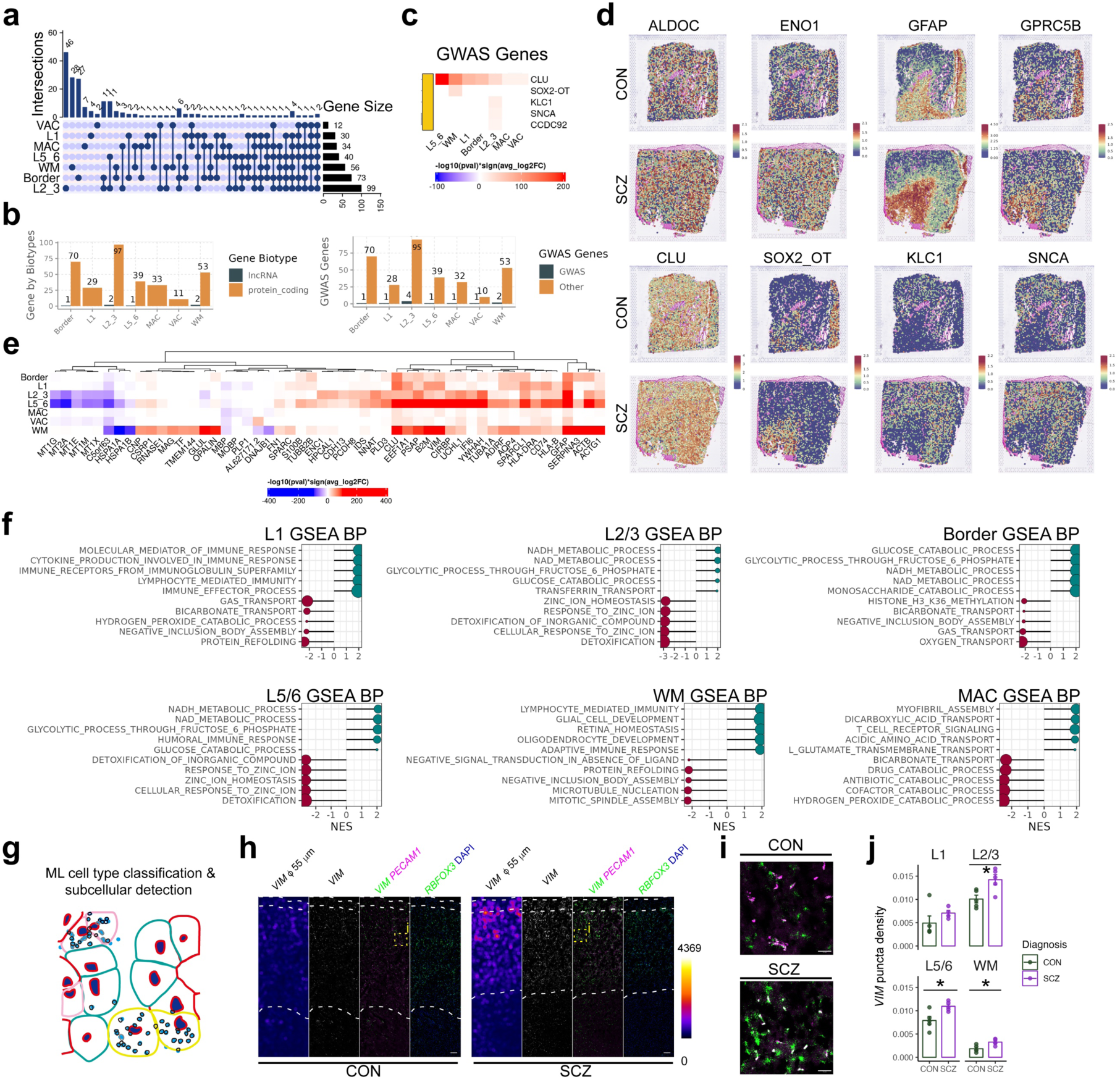
Spatial transcriptomic and pathway signature changes in the ACC in SCZ. **a**, Upset plot delineating the intersection of DEGs across five cortical spatial domains in SCZ. The size of the intersecting sets is represented by the connected dots, with the gene count corresponding to the bar length (*p*_adj_ < 0.05, FC > 1.2). **b**, Gene biotypes within identified DGE signatures from spatial domains in SCZ, showing the number of long non-coding and protein-coding genes (left panel) and the number of GWAS-associated genes (right panel). **c**, Heatmap depicting the identified GWAS genes associated with SCZ. **d**, Representative spatial expression of relevant genes and GWAS-associated genes in CON and SCZ cortical domains. **e**, Heatmap of top dysregulated genes (*p*_adj_ < 0.05, FC > 1.3) across cortical domains in SCZ, with hierarchical clustering illustrating patterns of dysregulation. **f**, GSEA displays biological process involvement in different cortical layers and domains in SCZ, with each domain’s key pathways annotated. The plot reveals domain-specific pathway alterations. **g**, Histological quantification of *VIM* expression. **h-i**, Representative RNA-FISH images. Average *VIM* intensity in 55 μm mimics Visium ST. Scale bars, 200 μm (**h**), 50 μm (**i**). **j**, *VIM* puncta density (/μm^2^, 5 individuals for each group [*df* = 8] except for L1 [5 CON and 4 SCZ, *df* = 7]. L1, *t* = 1.18, *p* = 0.27; L2/3, *t* = 3.03, *p* = 0.016; L5/6, *t* = 3.08, *p* = 0.015; WM, *t* = 3.19, *p* = 0.013, two-sided Student’s *t*-test). Mean ± SEM. \**p* < 0.05.

Numerous genes have been previously associated with SCZ in genome-wide association studies (GWAS). To identify DEGs genetically implicated in SCZ, we cross-referenced them to the genes reported in a recent study that focuses on fine-mapping and functional genomic data^21^. We identified five DEGs associated to SCZ in ST dataset. These genes include *CLU*, *SOX2-OT*, *KLC1*, *SNCA*, and *CCDC92*. *CLU* was upregulated across multiple cortical domains (Figs. 2c, and d, lower left panel), consistent with previous transcriptomics studies^3,22,23^. Astrocytes primarily secrete CLU to enhance excitatory synaptic transmission^24^; moreover, *CLU* represents an upregulated stress response gene in aging and neurodegeneration^25^. Intriguingly, our study also highlights the upregulation of the GWAS-linked long non-coding RNA (lncRNA) *SOX2-OT*, predominantly in the WM of SCZ patients. *SOX2-OT* expression has been reported to be altered in the olfactory epithelium of SCZ patients^26^. Sox2ot downregulates Sox2^27^, and Sox2 depletion leads to abnormal astrocyte morphology and physiology and impaired glutamate uptake in mice^28^. Our examination extended to other lncRNAs potentially linked to SCZ, though we observed minimal changes across ACC cortical layers at the tissue level (Fig. 2b, left panel).

To investigate the transcriptomic alterations deeper, we conducted Gene Set Enrichment Analysis (GSEA), revealing an enrichment of immune response genes in the L1, and enrichment in glial cell development in WM, alongside prominent metabolic pathway alterations involving NADH and glucose metabolism in L2/3, L5/6, and Border regions (Fig. 2f, Table 2). These observations highlight a strong influence of glial cells on the spatially resolved transcriptomic changes found in SCZ and suggest that glial cells may modulate the cortical microenvironment even in the layers where neurons predominate.

### Multiplex RNA-FISH confirmed astrocyte transcriptomic alterations

Among other glial-related genes, ST analysis highlighted dysregulated expression in astrocytic genes, such as *VIM* and *SERPINA3*. To confirm these findings, we performed RNA-FISH in five CON and five SCZ samples, including all the eight cases analyzed by Visium ST (Extended Data Table.1). We first investigated *VIM* as one of the possible astrocyte marker genes altered in SCZ. Because *VIM* is also expressed in endothelial cells, we co-stained with *PECAM1*, an endothelial marker. *RBFOX3*, a neuronal marker, was used to identify neurons and the GM. Using an unbiased imaging approach with machine learning-based cell classification (Fig. 2g; Methods), we classified more than 300,000 cells into *VIM*+ astrocytes (*VIM*+, *PECAM1*-, *RBFOX3*-), *VIM*+ endothelial cells (*VIM*+, *PECAM1*+, *RBFOX3*-), *VIM*-endothelial cells (*VIM*-, *PECAM1*+, *RBFOX3*-), neurons (*VIM*-, *PECAM1*-, *RBFOX3*+), and other cells (*VIM*-, *PECAM1*-, *RBFOX3*-; e.g. oligodendrocytes) (Figs. 2h-i). Notably, *VIM* puncta were enriched in *VIM*+ astrocytes and *VIM*+ endothelial cells (Extended Data Fig. 2a). It is well known that neurons, astrocytes, and oligodendrocytes have large, intermediate, and small (lymphocyte-like) nuclear size, respectively. The size of nuclei for each cell type classified was as expected, validating the cell classification (Extended Data Fig. 2b). As observed in ST and sn-RNA-seq data (Fig. 2e), *VIM* expression was markedly elevated across samples in the L2/3, in addition to the L5/6 and WM in SCZ (Figs. 2h-j, Extended Data Figs. 2c-d). We found a similar trend for increased cell density of *VIM*+ astrocytes but not *VIM*+ endothelial cells (Extended Data Figs. 2f, g). We further found a subtle but significant increase in the number of *VIM* mRNA molecules in each *VIM*+ astrocyte (from the L1 to WM) and *VIM*+ endothelial cell (from the L2-6) (Extended Data Fig. 2e), suggesting that the increase in *VIM* expression in SCZ observed in ST is mainly attributable to an increase in the number of *VIM*+ astrocytes. We also stained *SERPINA3*, with *RBFOX3* and *MBP*. The number of *SERPINA3* mRNA molecules in each *SERPINA3*+ astrocyte was significantly higher in SCZ in the WM (Extended Data Fig. 2h). The *SERPINA3*+ cell density and expression of *SERPINA3* average across all cells in each layer trended higher in SCZ but not statistically significant (Extended Data Figs. 2i, j), with one patient showing exceptionally high values. This observation was consistent with a previous study, in which high *SERPINA3* expression was observed only in a subgroup of SCZ patients with high inflammation^29^. Our RNA-FISH confirms the significant transcriptomic alterations related to astrocytes in both the WM and GM in SCZ and highlights spatial and donor-specific differences.

### Single-cell and spatial transcriptomics integration show regional-dependent glial alterations in SCZ

We found glia-specific alterations in our ST analysis that could be related to changes in cell type compositions. In this respect, contradicting results have been reported about alterations in neuronal and glial cell density in the cerebral cortex of SCZ patients^15,30,31^. Utilizing our sn-RNA-seq data generated from ACC, we annotated Visium ST spots to identify predominant cell type signatures (Fig. 3a; Methods). We then compared these signatures between CON and SCZ subjects. Our analysis revealed a pronounced upregulation of astrocytic signals, most notably in the Border and L1. Concurrently, we observed a modest increase in microglial signals within the GM and WM region in SCZ, contrasting with a decrease in oligodendrocyte signatures across the WM and GM. Excitatory neuron signatures presented a mixed pattern, diminishing at the Border while increasing in other cortical areas (Fig. 3b). To elucidate how similar the transcriptomic changes observed in the ST of SCZ are to the transcriptomic changes of a specific cell type, we pre-selected the top 100 DEGs in SCZ from our single-nuclei analyses to build a “SCZ cell-type specific response” custom gene sets for a GSEA on Visium ST data. This approach allowed us to determine a high similarity in L2/3 and L5/6 transcriptomic response to astrocytes and microglia (Fig. 3c). Additionally, the WM and Border regions showed notable similarities with oligodendrocyte precursor cells (OPCs) and oligodendrocytes, underscoring the pivotal role of glial cells in the observed transcriptomic alterations.

**Fig. 3.**
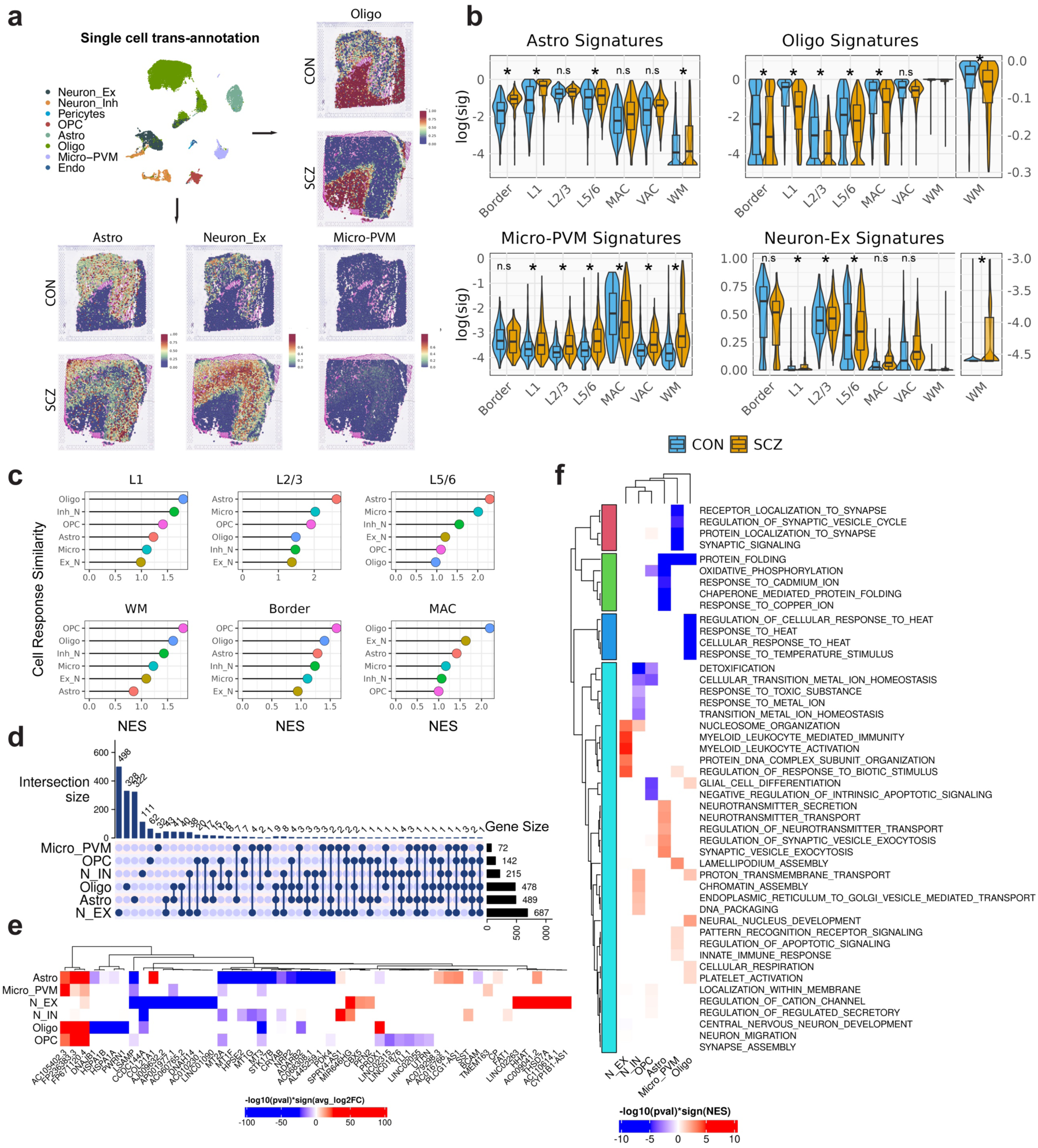
Cell Type-Specific Transcriptomic Alterations in SCZ. **a**, Single-cell trans-annotation of ST spots in the ACC depicting the spatial distribution of cell types in CON and SCZ subjects. Specific cell types are color-coded. **b**, Violin plots indicating the distribution of cell type-specific signatures in CON versus SCZ samples across cortical layers and regions. (log(signatures values of each Visium ST spot), two-way ANOVA followed by Tukey’s range test. Astro; Layers:Diagnosis, *F* = 87.03, *p* < 0.0001. Oligo; Layers:Diagnosis, *F* = 44.6, *p* < 0.0001. Micro-PVM;Layers:Diagnosis, *F* = 18.81, *p* < 0.0001; Neuron-Ex; Layers:Diagnosis, *F* = 29.58, *p* < 0.0001), \**p* < 0.05, n.s, no significance. **c**, Enrichment plots displaying cell type-specific similarity using custom GSEA Normalized Enrichment Scores (NES) in various cortical layers and regions. The color corresponds to the cell type. **d**, Upset plot delineating the intersection of DEGs across distinct cell type in SCZ. The size of the intersecting sets is represented by the connected dots, with the gene count corresponding to the bar length (*p*_adj_ < 0.05, FC > 1.2). **e**, Heatmap of top dysregulated genes (*p*_adj_ < 0.001, FC > 3.5) across ACC cell types in SCZ, with hierarchical clustering illustrating patterns of upregulation or downregulation. **f**, Heatmap with hierarchical clustering showing top 5 enriched and depleted GO, GSEA results in SCZ. The color gradient represents the –log10(*p*-value) * sign(NES) for enriched pathways across cell types.

Since Visium ST spots may include several cell types, we leveraged sn-RNA-seq data from the same tissue block to investigate cell-type specific gene expression changes and pathways altered in SCZ (Figs. 3d, e, f, Extended Data Figs. 3b and Table 3). We found the largest dysregulated genes in excitatory neurons with 687 genes, followed by astrocytes and oligodendrocytes with 487 and 478 genes, respectively (*p*_adj_ < 0.05 and FC >1.2). To highlight the most significant genes dysregulated in SCZ, we filtered DEG with *p*_adj_ < 0.0001 and FC > 3.5 (Fig. 3b). In excitatory neurons, we found that the limbic system-associated membrane protein (*LSAMP*) is the most downregulated gene (Fig. 3e and Extended Data Fig. 3c). *LSAMP* encodes a neural adhesion protein involved in neurite formation and outgrowth and in the integrity of serotonergic synapses^32,33^ and has been implicated in the etiology of SCZ^34,35^. In addition, we report the significant upregulation of the *HHAT* gene that encodes an enzyme responsible for N-terminal palmitoylation of Sonic Hedgehog (SHH) proteins involved in brain development^36^. In astrocytes, notable downregulations included *HPSE2*, *NRP2*, and *PDK4*, alongside the *MT* gene family—a finding echoed in spatial transcriptomics. Conversely, *COL21A1* and *CP*, and lncRNAs *AC105402.3 FP236383.3* and *FP671120.4* were upregulated (Fig. 3e and Extended Data Fig. 3c). Dysregulated genes associated with SCZ GWAS in ST (Fig. 2c) were also found in our DGE GWAS-associated sn-RNA-seq (Extended Data Fig. 3d). Interestingly, *CLU* and *SERPINA3*, both upregulated in the Visium ST data (Fig. 2e), exhibited divergent dysregulation patterns between single-nuclear and ST data, with *CLU* being downregulated and *SERPINA3* upregulated in the single-nuclear data (Extended Data Figs. 3c, d). This might be due to changes in the neuronal *CLU* expression in SCZ^22,23^, which was not observed in our sn-RNA-seq data, and possible differences in the *CLU* RNA localized in the nuclei versus cytoplasm. Oligodendrocytes displayed significant upregulation in *PLP1*, *OPALIN*, *CNP* and *RNASE1* (Extended Data Fig. 3c), which were observed dysregulated in spatial analyses (Fig. 2e). Downregulated genes, including *HSPA1A*, *PWRN1*, and *HSPA1B*, are often previously known to be associated with SCZ^37–39^ (Fig. 3e).

Further, GSEA revealed alterations in immune response pathways in excitatory neurons (Fig. 3f). Microglia showed dysregulation in pathways critical for synaptic signaling modulation and innate immune response. In OPCs, the observed downregulation of cell differentiation pathways (Fig. 3f, Table 4) may suggest diminished oligodendrocyte signaling in the L1, GM, and WM (Fig. 3b). Notably, astrocytes exhibited a dysregulation in oxidative phosphorylation of ATP synthesis—primarily driven by nuclear-encoded genes, and upregulation in neurotransmitter transport modulation. Taking all together, our single-cell and spatial transcriptomics integration revealed compositional changes of glial signatures, with the dysregulation of astrocytes, and identified novel and previously reported molecular alterations associated with schizophrenia, hinting at intricate crosstalk between neurons and different glial cell types related to immune response, differentiation, synaptic modulations, and metabolism.

### Immunohistochemistry for GFAP in a larger cohort reveals SCZ-specific morphological changes in a subset of astrocytes

Given the observation of astrocyte dysregulation at the molecular levels in our integrative spatial and single-nuclear analysis in SCZ, we performed IHC analysis of astrocytes in a larger cohort – 13 CON and 17 SCZ cases including all aforementioned samples – using formalin-fixed paraffin-embedded (FFPE) sections for better morphological preservation (Extended Data Figs. 4a-c, Extended Data Table 1). We performed IHC for a canonical astrocytic marker GFAP that allows the investigation of astrocytes of different subpopulations in the human cortex, including interlaminar astrocytes (ILA), protoplasmic astrocytes, varicose projection astrocytes, and fibrous astrocytes, which distribute in different cortical layers^40^ (Figs. 4a, b). Following our transcriptome findings in SCZ (Figs 2 and 3), we investigated whether the expression changes of astrocyte genes could indicate an alteration in the density of the astrocytic subtype population reported in the human cortex. First, we investigated the primate-specific ILAs characterized by having their soma localized in the L1 of the cortex and extending long processes to deeper layers of the cortex^41^. As expected, we found that ILA in CON samples exhibit long processes that extend from the pial and subpial surface in the L1 deep into the GM. Remarkably, in contrast, SCZ subjects had fewer ILA processes reaching the GM 100 μm deep from the L1 (light blue circles in the schema of Fig. 4a), indicating decreased abundance of ILA processes reaching the L2/3 (Figs. 4a-d, Extended Data Fig. 4d). We next focused on GFAP+ cells within the GM (hereafter GM refers to L2-6) that include protoplasmic and varicose projecting astrocytes, and found no statistical significance (Fig. 4d, middle panel). However, the density of GFAP+ cells in the WM was significantly higher in SCZ (Fig. 4d, right panel). Previous reports have shown sex differences in the protein content of GFAP^42^. We investigated the influence of sex on the GFAP+ differences found in SCZ and CON. We found a larger difference in male subjects with a significant increase in the L1 subpial GFAP+ cell density and WM (Fig. 4e). Moreover, the density of GFAP+ cells in the GM was higher in SCZ than CON in males with a modest significance (9 CON and 10 SCZ males, *p* = 0.065, two-sided exact Wilcoxon rank-sum test) (Fig. 4e). Similar trends were observed in the average GFAP intensity (Extended Data Figs. 4e-j), and percent (%) area positive for GFAP in the GM (Extended Data Figs. 4k, l).

**Fig. 4.**
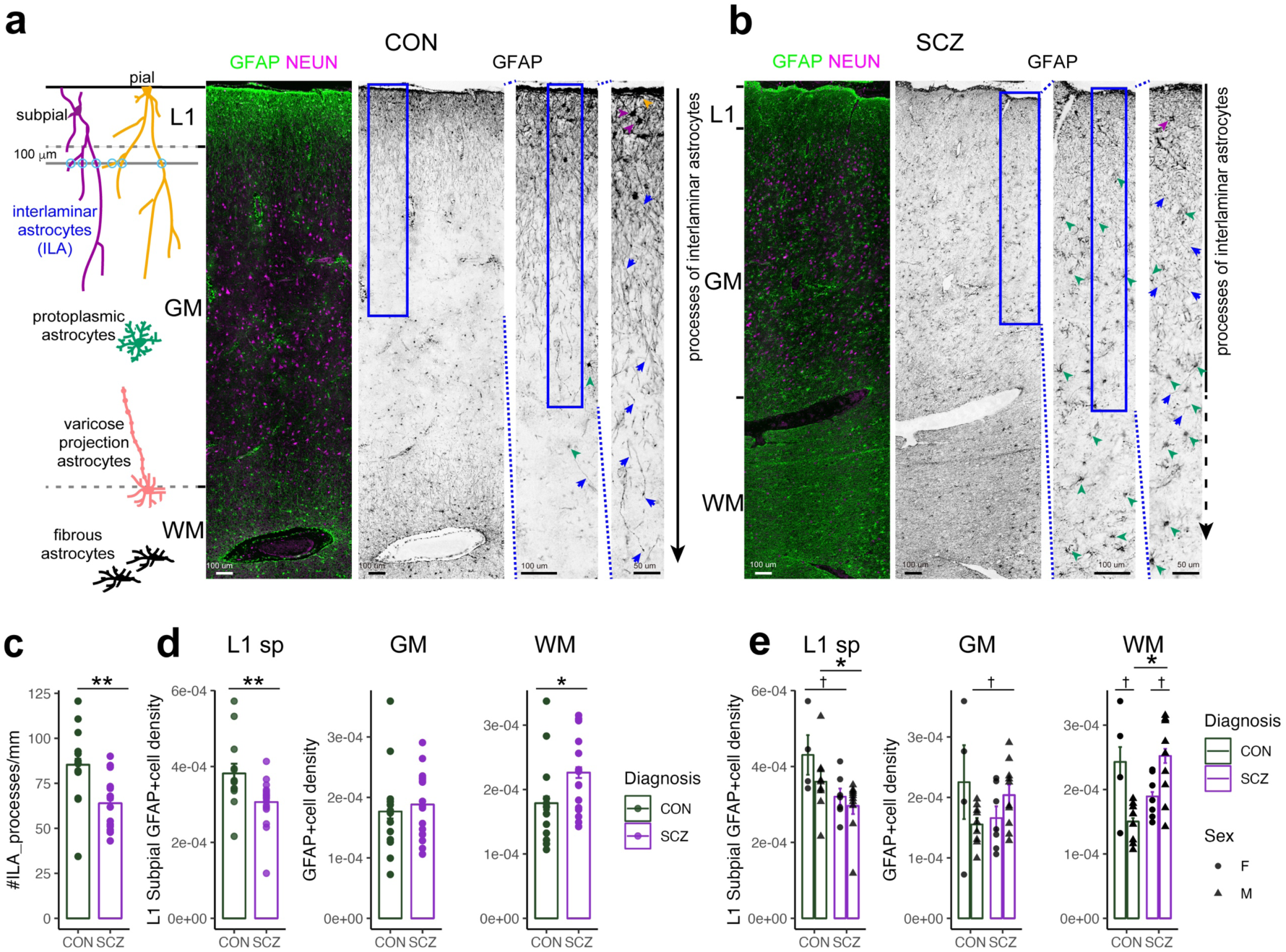
Primate-specific interlaminar astrocytes decreased in SCZ. **a-b**, Diagram explaining astrocyte heterogeneity (left panel) and representative image of the ACC stained for GFAP (right panels). The somata of ILAs are adjacent to the pial surface (orange arrowheads) or in the subpial L1 (purple arrowheads), extending long processes (blue arrows). Protoplasmic astrocytes are in the GM (green arrowheads showing small GFAP+ cells). In the deeper part are varicose projection and fibrous astrocytes. Scale bars, 100 and 50 μm, as indicated. **c**, The density of processes of ILAs at 100 μm from the L1/2 boundary (13 CON and 17 SCZ cases, *p* = 0.0045, two-sided exact Wilcoxon rank-sum test). **d**, The density of GFAP+ cells (/μm^2^) in the L1sp (L1 zone excluding the glial limitans), GM, and WM (13 CON and 17 SCZ cases; L1, *p* = 0.0045; GM, *p* = 0.46; WM, *p* = 0.039, two-sided exact Wilcoxon). **e**, Comparison by sex about GFAP+ cell density (/μm^2^) in the L1sp, GM, and WM (9 CON and 10 SCZ males; L1sp, *p* = 0.028; GM, *p* = 0.065; WM, *p* = 0.0021. 4 CON and 7 SCZ females, L1sp, *p* = 0.073; GM, *p* = 0.53; WM, *p* = 0.41, two-sided exact Wilcoxon). Sex differences within CON or SCZ were modestly significant in the WM GFAP+ cell density (CON, 4 females and 9 males, *p* = 0.076; SCZ, 7 females and 10 males, *p* = 0.055; two-sided exact Wilcoxon). * *p* < 0.05, ** < 0.01, † < 0.1. Mean ± SEM.

### Astrocyte alterations associated with aging and disease duration

Several lines of evidence show increased brain GFAP expression in aging^43^. To further elucidate the influence of aging on astrocytes, we performed a detailed histological examination of GFAP+ astrocytes across cortical layers in both CON and SCZ subjects (Figs. 5a, b). Although we found no linear association with aging in the abundance of ILA processes in CON and SCZ (Fig. 5c, upper panel), the GFAP+ cell density in the GM increased with a positive and significant correlation in CON (*ρ* = 0.95, *p* = 0.0001, Spearman correlation) (Fig. 5c, middle panel; see also Extended Data Figs. 5a-c). Furthermore, we noticed an inverse shift in the density of ILA processes and the density of GFAP+ cells in the GM in CON donors; that is, in many individuals, as the density of ILA processes decreased with aging, GFAP+ cell density increased. To capture this shift, we calculated an ILA index by dividing the density of ILA processes by the density of GFAP+ cells per individual. We found a significant, negative linear correlation between the ILA index and aging (*ρ* = –0.63, *p* = 0.02, Spearman correlation) (Fig. 5c, lower panel). Both ILAs and protoplasmic astrocytes could interact with neurons and blood vessels^44^ and, to a certain extent, show similar electrophysiological properties^45^, raising the possibility that they have partially overlapping functions. In SCZ, the ILA index was low, irrespective of age, resembling the astrocyte morphological patterns found in the elderly. In postmortem studies, disease duration indicates the time patients survived after their disease onset (Fig. 5e). We found a positive association (*ρ* = 0.57, *p* = 0.021, Spearman correlation) of the density of the ILA processes in SCZ, but not of the GFAP+ cell density in the subpial L1, GM, and WM, in case of the disease duration (Figs. 5g, h), suggesting some protective roles of the processes of the ILAs in SCZ. When comparing astrocytic parameters of GFAP IHC and clinical characteristics, we did not find significant associations, including age at onset, postmortem interval (PMI), disease severity as assessed by Diagnostic Instrument for Brain Studies (DIBS), and antemortem antipsychotic medication as assessed by chlorpromazine equivalent doses (Extended Data Fig. 5d).

**Fig. 5.**
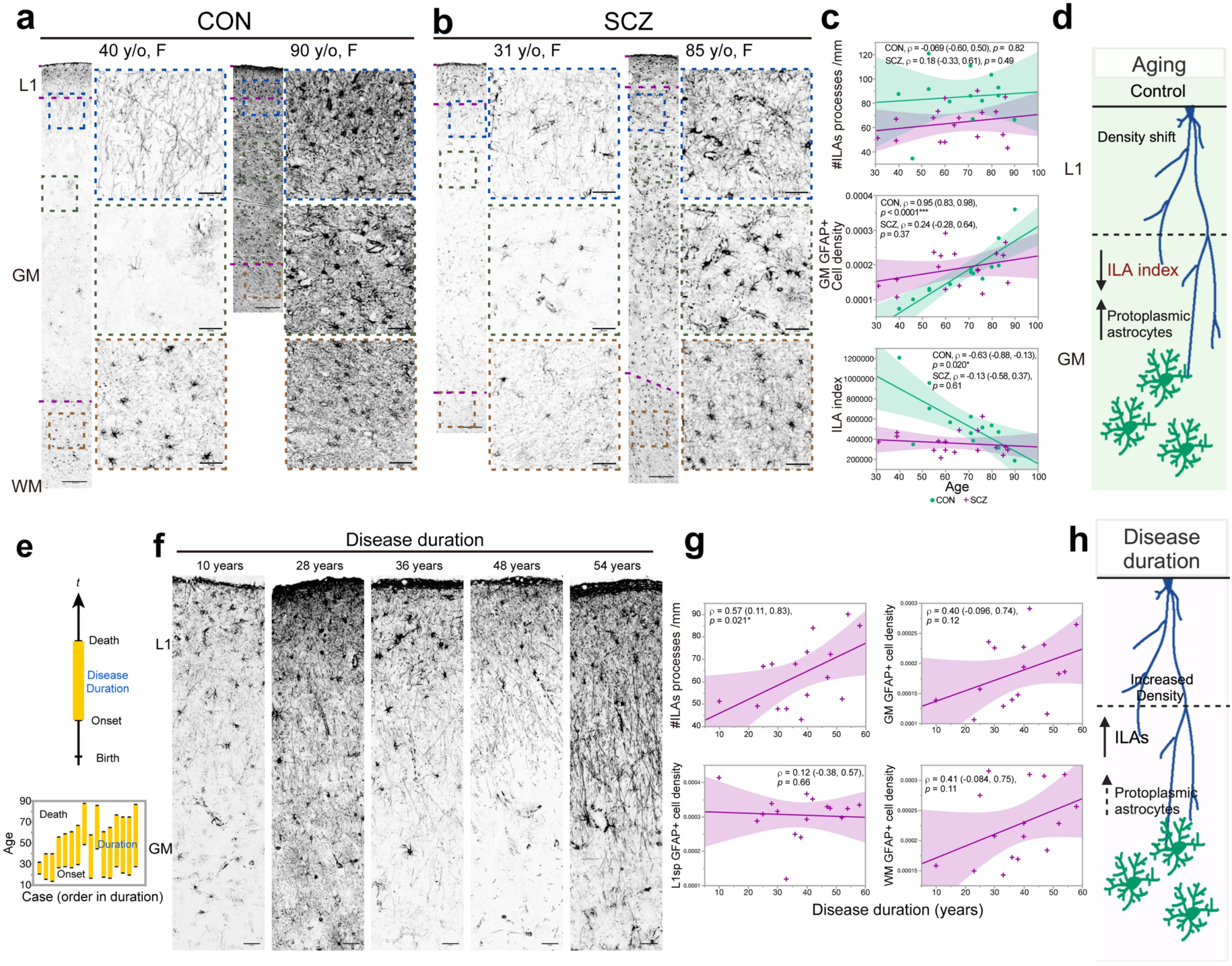
Microenvironment-specific GFAP + astrocytes changes in aging and disease duration. **a-b**, Representative immunohistochemistry images of GFAP from CON (**a**) and SCZ (**b**) donors of different ages (13 CON and 17 SCZ cases), displaying ILA process density and GFAP+ cell distribution, with age and sex indicated for each image. Scale bars, 200 μm, 50 μm. **c**, Relationship between age and the density of ILA processes, the density of GFAP+ cell in the GM, and ILA index, which indicates relative abundance of the processes of ILAs to GFAP+ cells in the GM, across CON and SCZ subjects (13 CON and 17 SCZ cases). The parameters of Spearman correlation [*ρ* (95% confidence interval), *p* value] are shown in the graphs. **d**, Schema depicting density shift in the GM from processes of ILAs to protoplasmic astrocytes. **e**, Schema depicting the relationship between aging and disease duration in postmortem brain research. Disease duration represents how long patients survived with the disease. **f**, Representative immunohistochemistry images of GFAP from SCZ patients with different disease duration (16 SCZ cases), displaying ILA process density. Scale bars, 50 μm. **g**, Positive association between disease duration and density of ILA processes (16 SCZ samples, *ρ* = 0.57 (0.11, 0.83), *p* = 0.021, Spearman) contrasts with nonsignificant associations of disease duration and the density of GFAP+ cell density in the L1sp, GM, and WM. **h**, Schema depicting characteristics of astrocytes associated with longer disease duration. * *p* < 0.05. Linear regression lines with a 95% confidence interval are shown.

### Molecular changes in the ACC communication networks in SCZ

Given the discovery of morphological and molecular changes in interlaminar astrocytes in SCZ, we aimed to interrogate the intercellular communication network with a spatial context using CellChat (v2)^46^. CellChat models the probability of interactions by integrating gene expression data and established interactions among signaling ligands, receptors, and cofactors. It employs the law of mass action^47^ to calculate communication probabilities. We found an increased number of overall predicted interactions in SCZ in all layer domains of the cortex, with a great extent in the L1 and WM of the cortex (Figs. 6a, b). In addition, the ligand-receptor (L-R) pairs detected in CON and SCZ samples are part of 29 signaling pathways, including the CypA (cyclophilin A or peptidylprolyl isomerase A, a gene product of *PPIA*), PSAP (prosaposin, involved in the hydrolysis of sphingolipids), PTN (pleiotrophin), MIF (macrophage migration inhibitory factor), SPP1 (secreted phosphoprotein 1), and FGF (fibroblast growth factor), among others (Fig. 6c). Interestingly, identified pathways such as CCK (cholecystokinin), IGFBP (insulin-like growth factor binding protein), CX3C (C-X3-C Motif Chemokine, or fractalkine), VISFATIN (nicotinamide phosphoribosyltransferase), EDN (endothelin), CSF (colony stimulating factor), PROS, EGF (epidermal growth factor), and CXCL (C-X-C motif chemokine ligand) were only found in SCZ. FGF, EGF, and BMP are critical in astrocyte differentiation and cell state changes^48–50^. The main predicted L-R contributors to these pathways showed a distinct spatial distribution pattern in the ACC cortical domains of SCZ donors. For example, the PTN, MIF, and SPP1 signaling increased in the L1 whereas the CypA signaling increased in L2/3, L5/6, and the WM (Figs. 6c-g and Extended Data Figs. 6b-c).

**Fig. 6.**
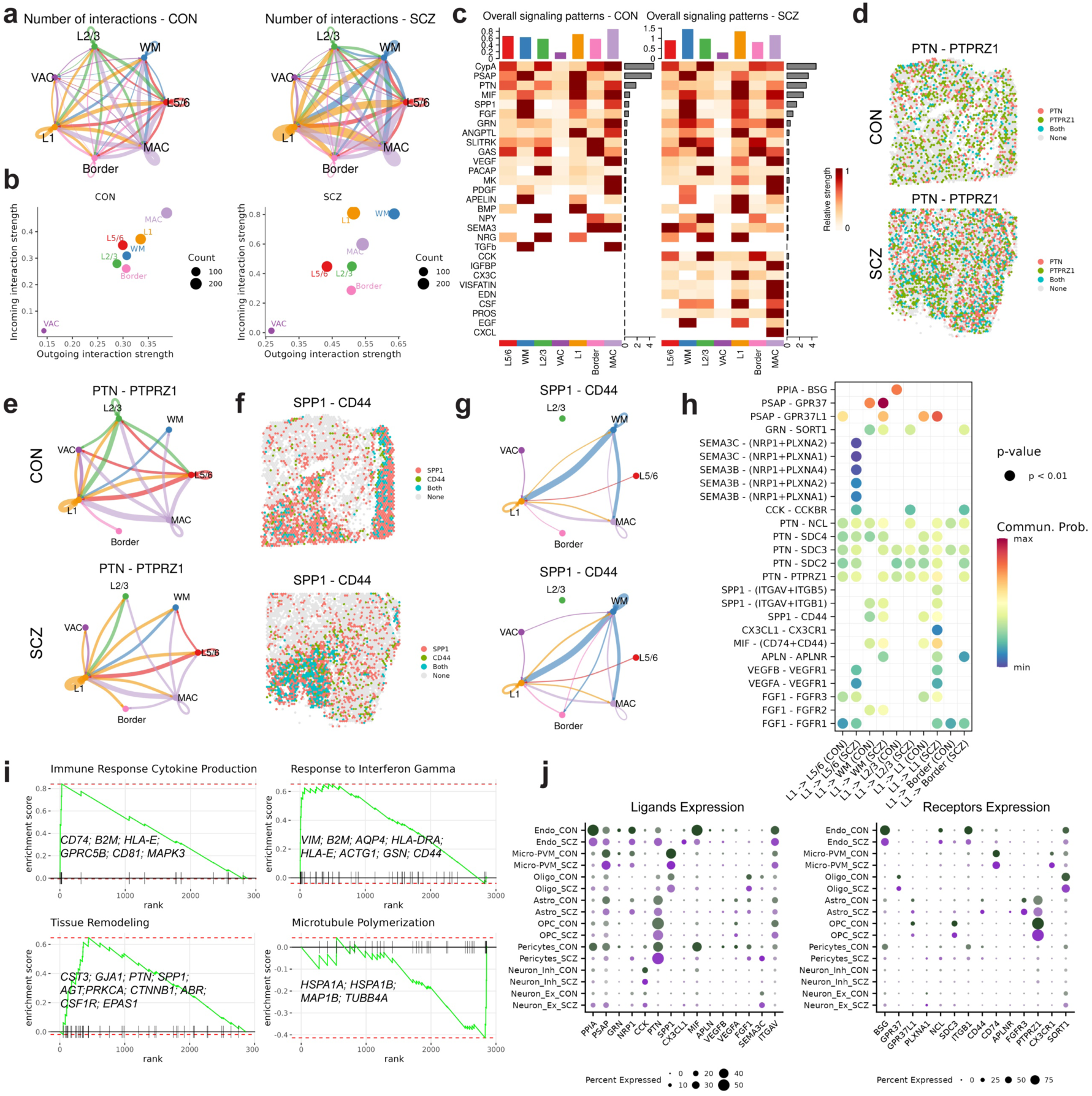
Predicted communication networks and signaling changes in the ACC L1 of SCZ donors. **a**, Number of cellular interactions in CON and SCZ conditions, depicted with network graphs for cortical domains-specific interactions. Each line of chord diagrams represents a different type of interaction or pathway, with the thickness indicating the strength of predicted communication. **b,** Scatter plots showing incoming versus outgoing interaction strength with dot size representing interaction count. **c**, Heatmaps comparing overall signaling patterns in CON and SCZ, with colors indicating relative signaling strength. **d**, Spatial distribution of PTN_PTPRZ1, main contributors of PTN signaling in CON and SCZ, illustrating alterations in signaling localization. **e**, Chord diagrams showing differences in predicted PTN regional communication. **f**, Spatial distribution of SPP1_CD44, main contributors to the SPP1 signaling in CON and SCZ, illustrating alterations in signaling localization. **g**, Chord diagrams showing differences in predicted SPP1 regional communication. **h**, L1 signaling diagrams indicating dysregulated L-R signaling interactions in SCZ, with dot colors denoting statistical significance (*p* < 0.01). **i**, Enrichment plots showing the involvement of L-R pairs in GSEA pathways, genes showing the genes that contribute most to the enrichment score (leading edge). **j**, Dot plots showing single-cell expression percentages of ligands and receptors in CON and SCZ.

We also performed an L-R pair statistical comparisons (Fig. 6h and Extended Fig. 6d), finding the most relevant predicted L-R pairs that include pathways modulated by NRP1, PTN, SPP1, PPIA, and CD74. Most of these ligands were lowly expressed in neuronal cells. Still, they do show the expression of their cognate receptors, indicating the potential modulation of neuronal signaling. Interestingly, pleiotrophin PTN is a signaling cytokine, highly expressed in astrocytes, OPC, oligodendrocytes, pericytes, and endothelial cells in our single-cell datasets (Fig. 6j). Astrocytic PTN reduces inflammation and decreases pro-inflammatory signals in both astrocytes and microglia and promotes neuronal survival^51^. SPP1, expressed by microglial cells and oligodendrocytes (Fig. 6j), is highly expressed in the L1, WM, and MAC and is predicted to interact with CD44 within and from these domains (Fig. 6h, Extended Data Fig. 6d). A previous report suggested that SPP1 has a neuroprotective role via the function of CD44^52^, which is expressed by astrocytes including ILAs^45^. It is noteworthy that polymorphisms in *PTN* are associated with the risk of SCZ^53,54^, and the levels of CD74 were reported to be downregulated in bulk tissue from the hippocampus and the amygdala in patients with SCZ, together with a non-significant upregulation in the ACC^55^.

Given our findings showing the lower abundance of subpial ILAs and their projecting processes in SCZ, we also investigated the involvement of identified L1 L-R in GSEA gene ontology (GO) pathways. We found the significant upregulation of immune response to cytokine production, response to interferon-gamma and tissue remodeling in L1, with *CD74*, *CD44,* and *SPP1* as genes that contribute the most to the enrichment score, respectively (Fig. 6i). Surprisingly, we also revealed a negative enrichment score for microtubule polymerization in L1 (Fig. 6i), which matches our IHC observations (Figs. 4a-c). The prevalence of CX3C signaling in the L1 of SCZ donors was a remarkable finding (Fig. 6h). The CX3CL1–CX3CR1 system plays a crucial role in brain homeostasis and microglia reactivity. While the CX3CL1 is present in the brain parenchyma and neurons, the CX3CR1 is mostly expressed in microglial cells^56,57^, and there are several lines of evidence showing the link of the CX3CR1 gene with SCZ at gene expression^58^ and even at the functional level, by linking the modulation of the CX3C signaling with SCZ-related behaviors^59^. On the other hand, CD74 and CD44 are predicted to interact with MIF in the L1 (Figs. 6h, j)^60^. CD74 is a marker for microglial activation, also known as major histocompatibility complex (MHC) class II invariant chain, which is highly expressed in microglial cells (Fig. 6j) and CD44 is highly expressed in astrocytes, indicating a possible microglial-astrocyte interaction as a major axis in the L1. This view is consistent with previous literature describing that activated microglia could drive neurotoxic astrocyte states^61^.

Taken together, our analysis suggests an intricate interplay between pro– and anti-inflammatory signals from microglial cells and astrocytes and raises the possibility that the changes in the ILA abundance could be a response to a pro-inflammatory microenvironment in the L1.

## Discussion

In this study, we employed an integrative approach that combines single-nucleus RNA sequencing, spatial transcriptomics, and histological analyses to explore the ACC region in patients with schizophrenia and control individuals. Our analysis has, for the first time, unveiled transcriptionally distinct regional domains within the ACC, shedding light on transcriptomic alterations in glial cells, predominantly in astrocytes. These findings led us to discover microenvironment-specific alterations in the composition and morphological complexity of a subset of primate-specific astrocytes, interlaminar astrocytes and found additional differences in sex and aging.

Our findings underscore the complexity of the cellular architecture in the ACC and its vulnerability in SCZ. Notably, the identification of microenvironment-specific alterations in the molecular composition and morphological complexity of ILAs marks a significant advance in our comprehension of the possible cellular mechanisms that may contribute to the pathology of SCZ. This finding challenges the traditional neuron-centric view of SCZ and suggests a critical role for astrocytes influencing the cortical microenvironment in the etiology of SCZ. Few reports have already highlighted astrocytic alterations in SCZ. A meta-analysis of transcriptomic studies covering five major psychiatric disorders found the astrocyte-related module up-regulated in bulk RNA-seq of SCZ brain samples^62^; more recently, neuron-astrocytic changes were characterized in a sn-RNA-seq study^5^. Several lines of genetic and morphological evidence suggest that SCZ is a disease with major alterations in the synaptic neuronal activity^21,63^. Since astrocytes are central to synaptic activity given their roles in glutamate recycling, D-serine NMDAR mediated stimulation, metabolic support, and facilitation of communication across neural and non-neural cells^64,65^, we propose that the dysregulation of astrocytes in SCZ, both molecular and morphological, may explain the neuronal dysfunction in SCZ. Although the role of ILAs is not well understood, ILAs display less abundance and processes in SCZ, which indicates fewer possible neuronal interactions and connectivity. In addition, our transcriptomic data indicates that in SCZ, astrocytes exhibit increased neurotransmitter release/transport and synaptic exocytosis. This is compelling since glutamate-induced Ca2+ transients in astrocytes, mediated by NMDA receptors, correlate with SCZ-like psychosis. Intriguingly, we observed reduced expression of metallothioneins, which are zinc-binding proteins with antioxidant properties, hypothesizing that local zinc dysregulation in the brain contributes to SCZ symptoms. Our findings also highlight metabolic alterations in SCZ, notably in NAD+/NADH metabolism and glycolysis, suggesting a redox imbalance in astrocytes. *in vivo* imaging^66^ and meta-analyses^67^ studies support this finding, indicating disrupted glutathione redox systems in SCZ, pointing to a potential therapeutic target in redox modulation. Additionally, our research suggests that inflammatory processes, as evidenced by astrocytosis and microgliosis, contribute to SCZ, potentially through cytokine-mediated neuronal damage.

A defining characteristic of the human cerebral cortex is the evolutionary thickening of the L2/3, distinguished by neurons exhibiting a rich diversity in transcriptomic profiles and possessing larger, more complex dendritic structures^68^. This feature highlights the intricate evolutionary path of cortical development, further evidenced by the substantial thickening of the subplate^69^, a region primarily diminishing postnatally but retaining some neurons in layers such as the L6b^70^ or WM. Intriguingly, the subplate is enriched with markers linked to SCZ and autism spectrum disorder^71^, exhibiting a potential vulnerability locus within the cortical architecture. Our analysis of Visium ST datasets reveals a region that we defined as the Border, a demarcation between the GM and WM, characterized by the expression of neuronal genes that could represent the L6b. Through this study, we observed a pronounced vulnerability of these two regions, by the large number of DEGs in the L2/3 and the Border, implicating these regions of evolutionary expansion of hominids as potential hotspots for schizophrenia susceptibility.

The ILAs distinctive to the L1 cortical layer in humans and primates exhibit extensive complexity and long projections not found in rodents or other higher-order mammals ^44^. Our data suggests that ILAs support cortical functioning by allowing better intracortical communication and neuronal metabolism, which declines with age and SCZ. ILAs could also be part of the glymphatic system since water channel AQP4-positive ILAs are associated with the glial limitans and cerebrovasculature, extending long processes from the cortical surface where branches of cerebral arteries distribute into the GM^72^. The shift from the ILAs to protoplasmic astrocytes in aging and SCZ (Figs. 4c, 5c-d) could result in alterations in such functions, given the striking differences in their morphology. We also showed that resilient individuals who survive longer with SCZ despite the multiple effects on their quality of life exhibit more ILA processes, suggesting positive roles of ILAs in maintaining individual health, given that people with schizophrenia die 10-15 years younger than the general population^73^. More investigation is necessary to understand the mechanisms by which ILAs modulate cortical function and the astrocytic dynamics in the ACC microenvironment and other regions of the human brain.

Our study faces several limitations: it is a case-control investigation, which cannot definitively establish causality with disease traits, highlighting the need for further animal or *in vitro* studies. Despite collecting information such as age, sex, postmortem interval (PMI), antipsychotic medication, and other known confounders in our analysis, the potential for unforeseen biases remains a common challenge in postmortem research. Although our sample size was relatively limited due to the challenges of obtaining high-quality fresh tissue of this particular brain region, we validated our transcriptomics findings through histological analysis in a larger cohort using fixed tissues. It remains to be determined whether the molecular and morphological findings are replicated in other brain regions. Further research is needed to establish the universality of our findings, considering the heterogeneity of SCZ and SCZ spectrum disorders across different cohorts worldwide. The spatial resolution of Visium ST technology, limited to 50 μm, precludes single-cell precision. We addressed this through a trans-annotation approach using single-nucleus RNA sequencing data from the matched sample source. Some genes exhibited divergent alteration patterns between single-nuclear and ST data, which can be due to changes in either cell population expressing the genes, or maturation of the mRNA causing different localization in the nuclei versus cytoplasm. Additionally, the morphological analysis of astrocytes was conducted on thin sections. We envision our future studies using three-dimensional high-resolution confocal imaging techniques to capture the cellular complexity of astrocytes and their interaction with other cell types.

In summary, our work illustrates the power of integrating diverse omics approaches to unravel the cellular and molecular complexity of poorly understood brain regions like the ACC in normal and in the context of SCZ. By shedding light on the role of astrocytes and highlighting novel cellular interactions, our study not only advances our understanding of SCZ neuropathology but also opens new avenues for therapeutic intervention. Future research should aim to further elucidate the functional implications of these findings, potentially paving the way for groundbreaking advancements in early diagnosis and treatment for SCZ.

## Acknowledgements

We would like to thank the donors of the brains and their families, whose gifts of hope made this study possible. We also thank 10X Genomics and ACGT Core for providing reagents and processing some of the samples. We are grateful to Tsukasa Kouno (RIKEN IMS), Ikuko Koya (RIKEN IMS), Shizuka Ohki (The Jikei University), and Maiko Saito (The Jikei University) for technical assistance; Hiromi Onuma (Tohoku Postmortem Brain and DNA Bank for Psychiatric Research and Tohoku University) for coordinating donations; Miho Ito (RIKEN IMS) for coordinating research collaboration. This work was supported by Grants-in-Aid for Scientific Research of the Ministry of Education, Culture, Sports, Science, and Technology (MEXT), Japan/Japan Society for the Promotion of Science, Japan, Grants-in-Aid for Scientific Research (KAKENHI) (JP22K15203, JP21H02853, JP21H00180), PRIME, AMED, Japan (JP19gm6310004, JP19dm0207074, JP21wm0425019), a grant from SENSHIN Medical Research Foundation, The Uehara Memorial Foundation, The Naito Foundation, Keio Gijuku Academic Development Funds, The Jikei University Exploratory Collaboration Research Fund, and The Jikei University Strategic Prioritized Research Fund. This publication is part of the Single Cell Medical Network of Japan and was also funded by a RIKEN IMS research grant from MEXT.

## Statements and Declarations

### Author Contributions

J.L., S.Y., K.K., and J.S. conceived the project and designed the research. K.K., J.S., Y.K., C.C.H, P.C and C.C. supervised the project. Y. A., A. K., K. H., and K. N. coordinated the project. M.H., A.N., and Y.K. collected samples and clinical information. J.L. performed all bioinformatic analyses and visualized the results. J.L., S.Y., M.H., K.K., and M.K. processed the tissue and performed experiments. M.J. performed data analysis. S.Y. performed histological examination and data analyses. J.L. and S.Y. led the project, interpreted the results, and wrote the manuscript with input from all co-authors. All authors approved the final manuscript.

### Materials & Correspondence

Jay W. Shin and Ken-ichiro Kubo

### Tables

Table 1. Summary_DGE_All_ACC_Domains_ST

Table 2. Summary_GSEA_GO_bp_All_ACC_Domains_ST

Table 3. Summary_DGE_All_Brain_Cells_sn-RNA-seq

Table 4. Summary_GSEA_GO_bp_All_Brain_Cells_sn-RNA-seq

## Methods

### Human postmortem brain tissue collection

Postmortem anterior cingulate cortex (ACC) samples from people with SCZ and CON subjects were obtained from the Tohoku Postmortem Brain and DNA Bank for Psychiatric Research. The present study was approved by the Ethics Committee of Fukushima Medical University (1685), Tohoku University Graduate School of Medicine (2021-1-127, 2022-1-213), RIKEN Yokohama Campus (H30-26), Keio University School of Medicine (2019-0212), and The Jikei University School of Medicine (33-438(11065)), and complied with the Declaration of Helsinki and its amendments. All procedures were carried out with the informed written consent of the next of kin. We used samples without a clinical history of neurodegenerative disease. Only one SCZ sample and one CON in this study had an early clinical history of cerebrovascular diseases, but macroscopic and nuclear staining assessment determined no evident related lesions in the area which we analyzed. In addition, 10 out of 31 samples were analyzed by board-certified neuropathologists, exhibiting no neurodegenerative or ischemic changes in the ACC, assessed by neuropathological staining such as hematoxylin/eosin and Klüver-Barrera staining (Nissl + Luxol Fast Blue). Demographic information on brain tissues is summarized in Extended Data Table 1 and Extended Data Figs. 4a-c (formalin-fixed, paraffin-embedded, or FFPE). Clinical diagnosis of the patients was performed based on the Diagnostic and Statistical Manual of Mental Disorders (DSM)-IV^74^, or 5^75^. The daily dosage of antipsychotics prescribed during the 3 months before death is shown as a chlorpromazine-equivalent dose in mg/day (CP-eq). The Diagnostic Instrument for Brain Studies (DIBS)^76^ was used to retrospectively estimate the symptoms of people with SCZ in 3 months before their death or most severe symptoms in life. We classified antemortem symptoms of DIBS into three subscales (positive symptoms, negative symptoms, and general psychopathology) and the total of these subscales.

### Sample processing

Fresh frozen tissue chunks containing both the gray and white matter of the dorsal ACC (6 CON and 6 SCZ) were trimmed to meet the size requirements for Visium spatial transcriptomics (ST), and cut coronally into two essentially identical, thin tissue blocks. One of the blocks was used to obtained nuclei and subjected to the 3’– and 5’-Single-nuclear RNA sequencing (sn-RNA-seq) (4 CON and 4 SCZ; 2 CON and 2 SCZ, respectively; see Extended Data Table. 1). The other block was embedded in the OCT compound (Sakura, Tokyo, Japan) and cryosectioned by 10 μm thick for Visium ST and histological analysis. FFPE samples were prepared from the contralateral hemisphere to the fresh frozen tissue fixed with 10% formalin at room temperature (RT) and embedded in paraffin according to the standard protocol. We prepared 7 μm thick sections using a microtome in 14 CON and 17 SCZ samples.

### sn-RNA-seq and Visium ST

Samples for 3’-sn-RNA-seq (Multiome kit) and Visium ST were processed by the AGCT Core (Los Angeles, USA), and IMS RIKEN for the generation of 5’-RNAseq following standard procedures—only the transcriptomic dataset was used from the multiome data. Briefly, cryopreserved tissues from the human ACC were mechanically dissociated using the Singulator device (S2 Genomics) to isolate nuclei following the manufacturer’s procedures. Prior to the protocol’s initiation, both lysis and washing buffers were prepared as previously described^77^. The lysis buffer was prepared with the following concentrations: 10 mM Tris-HCl (pH 7.4), 10 mM NaCl, 3 mM MgCl_2_.6H2O, and 0.05% NP-40 (v/v). The washing buffer contained 5% BSA (w/v), 0.25% glycerol (v/v), and 40 units/mL Protector RNase inhibitor, diluted in 0.5× PBS. Following tissue dissociation, the nuclei suspension was collected and filtered through a 20 µm cell strainer. Washing involved adding an equivalent volume of washing buffer to the filtered suspension and centrifuging at 400 g for 5 minutes at 4°C. The pellet was resuspended in fresh washing buffer, centrifuged at 200 g for 5 minutes, and passed through the strainer again. Nuclei were counted with LUNA-FX7™ Automated Cell Counter. Concentration of isolated nuclei was adjusted to target ∼10,000 nuclei. The nuclei were loaded onto the Chromium Controller (10x Genomics) using Chromium Single Cell 5′ Reagent Kit v1.1 and chip G (10x Genomics). RT reaction was performed at 53°C for 45 minutes according to manufacturer’s instructions. Then, cDNAs were amplified using cDNA primer mix in the kit, with 13 PCR cycles, followed by the standard steps in manufacturer’s instructions. The size profiles of libraries were examined by Bioanalyzer (Agilent) and the amount was quantified by KAPA Library Quantification Kits. These libraries were sequenced on MGIseq2000 (BGI): Read1 was 111 bp including dark cycle 27-39 and Read2 was 91 bp.

For visium ST, 10 μm-thick, fresh-frozen sections were adhered on gene expression slides in Visium Spatial Gene Expression Slide & Reagent Kit, 4rxns (10X GENOMICS, PN-1000187). After methanol fixation, hematoxylin and eosin staining was performed according to the manufacturer’s protocol (CG000160), and slides were imaged. Coverslips were removed and sections were processed using Visium Spatial Gene Expression Slides & Reagents (10X GENOMICS, PN-1000187). Sections were permeabilized for 18 minutes, based on results obtained from Tissue Optimization Slides (10X GENOMICS, PN-1000193). Reverse transcription, second strand synthesis and cDNA denaturation, cDNA amplification, library preparation, and sequencing according to the manufacturer’s protocol (CG-000239).

## Bioinformatics and Transcriptomic Data Analysis

### Single Cell RNA-seq Analysis

sn-RNA-seq reads were aligned to the human genome (hg38) using CellRanger version 3.1.0 (10x Genomics). The resulting Cell Ranger outputs were imported into R (version 4.12) for analysis with the Seurat package (version 4.1). Nuclei were filtered based on the following criteria: feature counts between 350 and 5,000, and mitochondrial gene percentage below 15%. Variance stabilization and normalization of individual datasets were performed using SCTransform^78^ (vst.flavor = “v2”), which enables downstream differential expression analysis. Integrated analysis was conducted with the Harmony package (version 1.2), facilitating data projection into principal components (PCs). Clusters were identified using a resolution of 0.8 and visualized using UMAP dimensional reduction with 30 Harmony-corrected PCs. Cell annotations were performed using the Azimuth package (version 0.46) using the human cortex reference and confirmed by known gene markers. For nuanced interpretation, neurons were categorized into excitatory and inhibitory groups. Prior to differential expression analysis, counts were recorrected using the PrepSCTFindMarkers function that uses a minimum of the median UMI (calculated using the raw UMI counts) of individual objects to reverse the individual SCT regression model using a minimum of median UMI as the sequencing depth covariate given multiple SCT models. We excluded mitochondrial and ribosomal genes in the analysis to avoid possible technical influences. Differential gene expression analysis was performed using a logistic regression (LR) having “project” (3’and 5’RNAseq) and “nCounts” as latent variables.

### Visium Spatial Transcriptomics Analysis

Visium spatial transcriptomic reads were aligned to the human genome (hg38) using SpaceRanger version 2.0.1 (10x Genomics). Data were processed in R (version 4.12) using the Seurat package (version 4.1), with nuclei filtering criteria set to feature counts between 350 and 5,000, mitochondrial gene percentage below 40%, and hemoglobin genes (hb) below 20%. SCTransform (vst.flavor = “v2”) was applied for variance stabilization and normalization. Data integration was achieved using Harmony (version 1.2). Clusters, derived from UMAP dimensional reduction of 30 Harmony-corrected PCs, were annotated based on spatial location and known gene markers. Similar to the single-cell differential gene expression analysis, we exclude mitochondrial and ribosomal genes in the analysis to avoid possible technical influences. PrepSCTFindMarkers was employed for count recorrection prior to utilizing a logistic regression model with “nCounts” as a covariate for DGE analysis. Spatial data were further analyzed using the Seurat trans annotation pipeline, with reference to our sn-RNA-seq dataset. We used CellChat (version 2.1) for ligand-receptor (L-R) interaction analysis with interaction.range = 300, contact.dependent = TRUE and contact.range = 100. Differentially expressed L-R pairs were filtered out when not present in at least three samples per group.

### Gene Set Enrichment Analysis (GSEA)

Gene set enrichment analysis for GO Biological Processes was conducted using the fgsea package (version 1.24) in R, with parameters set to minSize = 15, maxSize = 500, and nPermSimple = 10,000. Pathway analyses were based on gene ranks from differential expression analysis in both sn-RNA-seq and Visium spatial transcriptomics, including all compared genes. For cell type response similarity enrichment, top 100 dysregulated genes in SCZ corresponding to each determined cell type in sn-RNAseq were used to generate a custom gene panel for GSEA in Visium spatial DEG.

### RNA-fluorescence *in situ* hybridization (FISH)

We performed *in situ* hybridization on fresh frozen ACC samples (6 CON and 6 SCZ) using RNAscope^®^ Multiplex Fluorescent Reagent Kit v2 (ACDbio, 323110) following the manufacturer’s protocol with some modifications. Briefly, 10 μm sections on slide glass (Fisherbrand Superfrost Plus, Fisher Scientific, 12-550-15) from the deep freezer were equilibrated at room temperature for 10-30 min to recover from ice crystal formation and fixed with 4% paraformaldehyde for 60 min at RT. Washing twice with 1X PBS, slides were dehydrated with 50%, 70%, and 100% ethanol for 5 min at RT. After drying at RT for 5 min, slides were treated with hydrogen peroxide at RT for 10 min, followed by Protease IV treatment at RT for 30 min. The probes used in this study includes RNAscope® Probe-Hs-VIM-C2 (ACDbio, 310441-C2), Hs-RBFOX3(415591), Hs-PECAM1-O1-C3 (487381-C3), Hs-RBFOX3-C3 (415591-C3), Hs-MBP-C2 (411051-C2), and Hs-SERPINA3 (412671). C2 and C3 probes were diluted in C1 probes at a 1:50 ratio and incubated for 2 hr at 40 °C. After amplification, signals were amplified with TSA PLUS CYANINE 3 (AKOYA Biosciences, NEL744001KT), TSA PLUS CYANINE 3 (AKOYA Biosciences, NEL745001KT), and OPAL 520 (AKOYA Biosciences, FP1488001KT) with 1X Plus Amplification Diluent. To quench the autofluorescence, slides were incubated with 0.25X TrueBlack® Lipofuscin Autofluorescence Quencher (Biotium, 23007) for 30 sec at RT. 4’,6-Diamidino-2-phenylindole Dihydrochloride (DAPI, Nacalai, 11034-56) was added before mounting using PermaFluor Aqueous Mounting Medium (TA-030-FM, Thermo Fisher Scientific).

### Immunohistochemistry

The sections were deparaffinized with Xylene at RT for 5 min twice and rehydrated by incubating slides with 100% (5 min), 100% (5 min), 90% (3 min), 70% (3 min), 50% (3 min) ethanol, 1X PBS (3 min), and 1X PBS (3 min) at RT. Slides were then immersed in PBS with 0.01% Triton X-100 (Sigma-Aldrich) (PBS-Tx) for over 30 min at room temperature (RT). Antigen retrieval and blocking endogenous peroxidase activity were performed by incubating the sections in 1X AR6 buffer (AKOYA Biosciences, AR6001KT) at 105 °C for 15 min. The sections were incubated overnight with primary antibodies at 4 °C or 1 hr at RT. The primary antibodies used were rabbit polyclonal anti-GFAP (DAKO, IR524; ready-to-use), rabbit polyclonal anti-MBP (Proteintech, 10458-1-AP; 1:1000), rabbit polyclonal anti-NEUN (ABN78, Abcam; 1:800), and mouse monoclonal anti-NEUROFILAMENT (clone SMI31.1, BioLegend, 837801; 1:200). After three washes with PBS-Tx, the sections were incubated with horseradish peroxidase (GFAP and MBP)-, biotin (SMI31.1)-, or Alexa647-conjugated secondary antibodies raised in donkeys for 1 hour at RT. Sections were further incubated with streptavidin-horseradish peroxidase in the case of SMI31.1, followed by signal amplification with TSA PLUS CYANINE 3 (AKOYA Biosciences, NEL744001KT), TSA PLUS CYANINE 3 (AKOYA Biosciences, NEL745001KT), or OPAL 520 (AKOYA Biosciences, FP1488001KT) with 1X Plus Amplification Diluent. To perform multiplexed immunohistochemistry using antibodies raised in rabbits, staining of one of the antigens was performed, which was followed by antibody removal by incubating the sections in 1X AR6 buffer at 105 °C for 15 min, and then staining of another antigen was performed. We performed counterstaining with DAPI or DAPI + fluorescent Nissl (NeuroTrace™ 435/455, ThermoFisher Scientific, N21479). Sections were mounted using PermaFluor.

### Image acquisition and quantification of histological samples

Images were acquired using a confocal laser-scanning microscope (LSM880, Zeiss), or ALL-in-One fluorescence microscope (BZ-X810, Keyence) avoiding saturation of the pixels, and were stitched using ZEN (Zeiss) or BZ-X800 Analyzer (Keyence). Images were further analyzed using the Fiji software^79^ (Fiji/ImageJ2.14.0/1.54f). Maximum projection images of the tiled images were generated to evaluate all the cells in the gray matter (GM) and white matter (WM) in the regions examined. Machine learning-based automatic classification of cell types was performed using QuPath^80^(v0.4.3). We detected cells based on nuclear staining. We manually classified 300-700 cells/channel for training in one of the samples. We then combined the cell classifiers to classify cell types in all of the cell types in the layer (L) 1, L2-6 (GM), and WM. We described the cell type classification as follows (Fig. 2g and Extended Data Figs 2a-d): neurons, *VIM*-/*PECAM1*-/*RBFOX3*+; oligodendrocytes, *VIM*-/*PECAM1*-/*RBFOX3*-; *VIM*-endothelial cells, *VIM*-/*PECAM1*+/*RBFOX3*-; *VIM*+ astrocytes, *VIM*+/*PECAM1*-/*RBFOX3*-; *VIM*+ endothelial cells, *VIM*+/*PECAM1*+/*RBFOX3*-. Likewise, *SERPINA3*+ astrocytes refer to *SERPINA3*+/*RBFOX3*-/*MBP*-cells. *The subcellular detection* function of QuPath was also used to count the *VIM* and *SERPINA3* mRNA expression by counting the number of puncta. High grey cell coefficient ^81^ of the human cortex and scattered distribution of mRNA puncta made the visual interpretation of the abundance of transcripts at low magnification difficult.

To overcome this, we averaged signal intensities in nearby pixels at ø = 55 μm as indicated in Figs. 1d, 2h and Extended Data Figs. 2c and d, mimicking Visium. The GFAP-positive area was estimated by defining an appropriate threshold in immunohistochemical analysis. To evaluate the morphological parameters of GFAP-positive cells, we defined 2-3 (3 if applicable) rectangle regions of interest (ROIs) of 1 mm width at regions where the section is locally parallel to the processes of the ILA, and mean values from these ROIs were calculated to minimize the effects of section angle variation. The density of the processes of the ILAs was evaluated by counting the intersection of GFAP+ processes and a virtual line that is 100 μm deep from a boundary between the L1 and L2. The processes of ILAs, which usually extend from the L1 in a relatively parallel manner, were morphologically discriminated from processes of nearby protoplasmic astrocytes, which usually extend processes radially from their soma, and endfeet that cover the blood vessels, which resemble a tram track. We defined the ILA index as the density of the processes of the ILAs (*per* mm) / the density of GFAP+ cells in the GM (*per* μm^2^).

### Statistical Analysis and Visualization

Data analyses were performed using custom R scripts and JMP® Pro (version 16.2.0). Statistical significance between groups was determined using appropriate parametric and nonparametric tests, detailed in figure legends. Adjustments for multiple comparisons were made to control the familywise error rate, considering *P* or adjusted *P* values < 0.05 as statistically significant, and *P* value < 0.1 as modestly significant. To calculate the confidence interval of Spearman’s *ρ*, Langtest (Version 1.0) [Web application] from http://langtest.jp^82^. For data visualization, we used the ggplot2 (3.5.0), scCustomize (version 2.1), ComplexHeatmap (version 2.14), patchwork (version 1.1.2) packages in R, and JMP® Pro (version 16.2.0). The bar plots are shown with mean ± standard error of means (SEM), and box plots are shown with the center line (median), box limits (upper and lower quartiles), whiskers (The upper/lower whisker extends from the hinge to the largest/smallest value no further than 1.5 * interquartile range), and all the individual data points.

## Data and code availability

The datasets generated and analyzed during the current study are available in the Synapse repository, accessible via the following link: https://doi.org/10.7303/syn61253365. This repository includes all raw and processed data files and relevant metadata to facilitate reproducibility and further analysis. Users can access the data following registration and compliance with data-sharing regulations. All code is deposited in https://github.com/JulioLeonIncio/SCZ_ACC_L1_Astrocytes.

The imaging data that support the findings of the current study are available from the corresponding author (K.K.) upon reasonable request.

---

**Extended Data Fig. 1.**
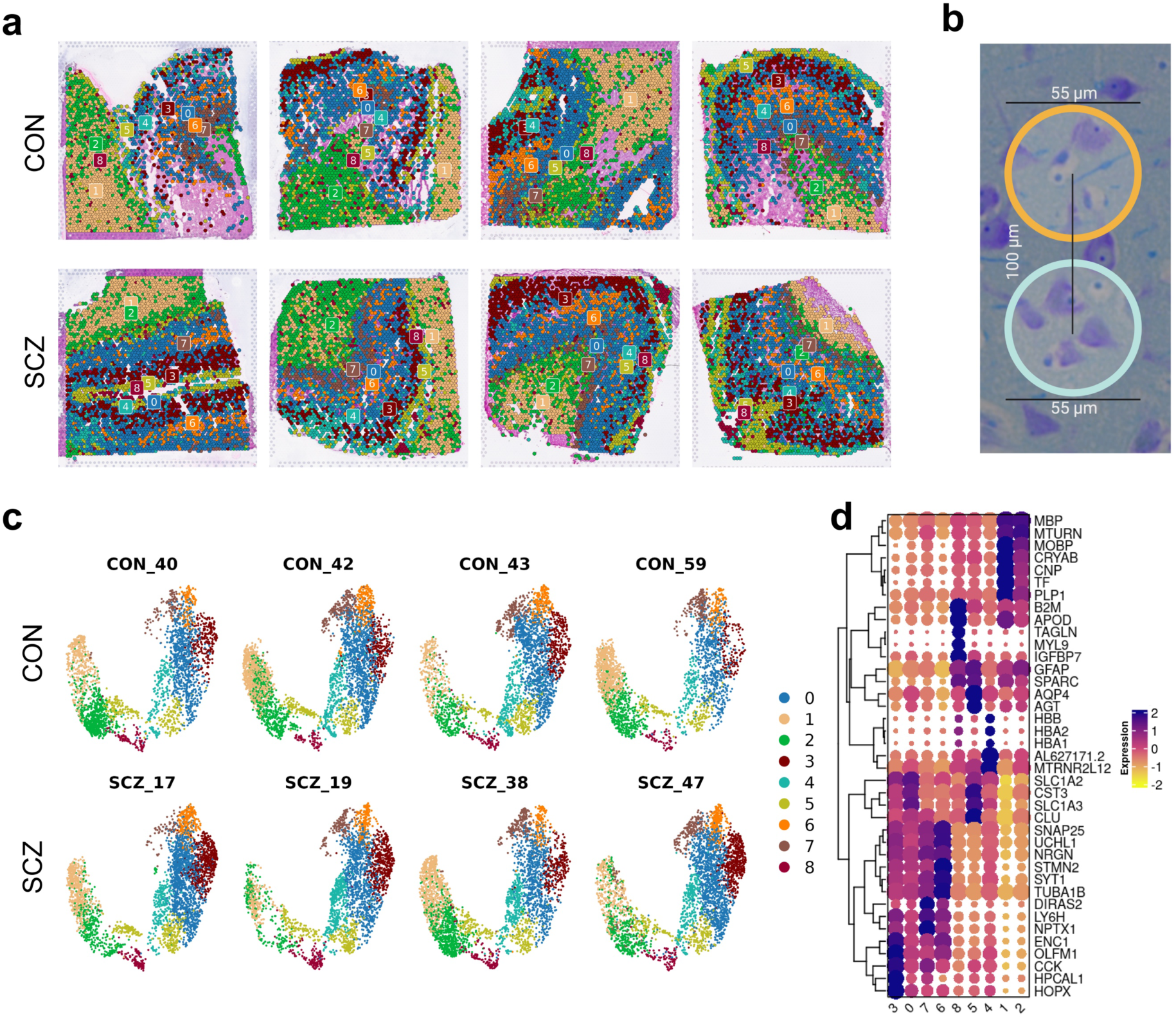
Identified ACC transcriptomic regional domains via unbiased clustering of Visium ST. **a**, Spatial clustering analysis of the ACC tissue sections. Top row: CON samples. Bottom row: SCZ samples. Each section is color-coded to represent distinct transcriptomic clusters identified through unbiased clustering. **b**, Illustration depicting Visium ST dots in the context of the cell composition and interactions in the human cortex —Nissl + Luxol Fast Blue staining. **c**, UMAP plots of spatial transcriptomics clusters. Individual plots correspond to CON donors 40, 42, 43, and 59, and SCZ donors 17, 19, 38, and 47. **d**, Heatmap of Marker Genes for Clusters Profiles.

**Extended Data Fig. 2.**
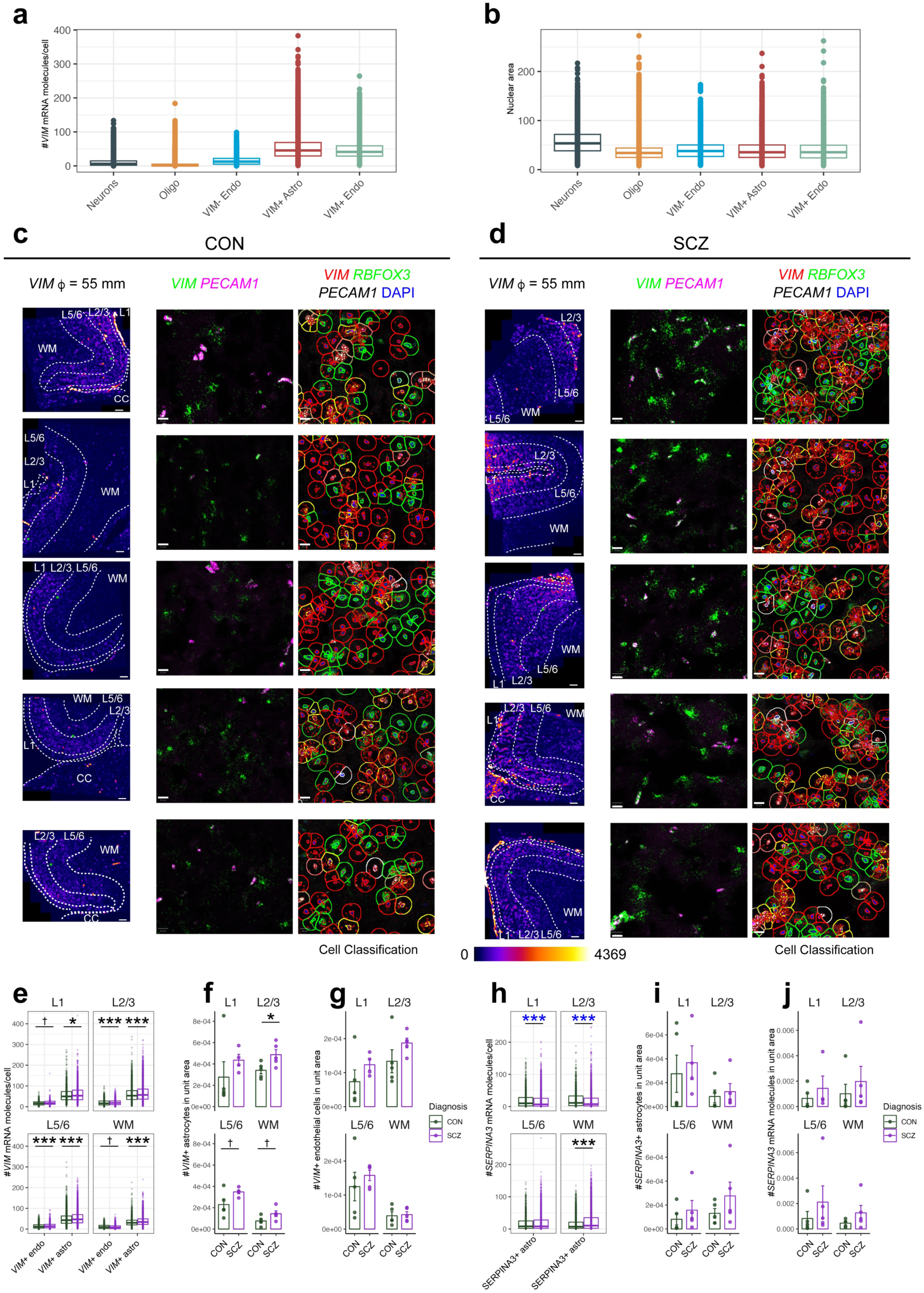
RNA-FISH astrocytes gene expression validation in the samples used for transcriptomic analysis. **a**, The number of *VIM* mRNA molecules/cell in different cell types. **b,** Nuclear size per cell type. **c-d**, All of the sections of CON (**c**) and SCZ (**d**) on which RNA-FISH for *VIM*, *RBFOX3*, and *PECAM1* was performed. High magnification is from the green asterisk in the low magnification, shown with automatic cell classification and *VIM* puncta detection. A fire look-up table of average intensity in a circle of 55 μm diameter is shown. Scale bars, 500 μm, and 20 μm. **e**, The numbers of *VIM* puncta in each *VIM*+ astrocyte (L1, *n* = 1258 cells [CON] and 2465 cells [SCZ], *p* = 0.010; L2/3, 4480 and 7532 cells, *p* < 0.0001; L5/6, 4542 and 6150 cells, *p* < 0.0001; WM, 2244 and 4813 cells, *p* < 0.0001, two-tailed Wilcoxon rank sum test; these cells are from 5 CON and 5 SCZ individuals except for L1 [5 CON and 4 SCZ], which is also applied for **h**) and *VIM*+ endothelial cells (L1, 341 and 726 cells, *p* = 0.060; L2/3, 1765 and 2992 cells, *p* < 0.0001; L5/6, 2156 and 2758 cells, *p* < 0.0001; WM, 1194 and 1330 cells, *p* = 0.091, two-tailed Wilcoxon). **f-g**, The cellular density (/μm^2^) of *VIM*+ astrocytes (*n* = 5 for each group except for L1 [5 CON and 4 SCZ samples]. This *n* is applied for **f**, **g**, **i**, and **j**; L1, *t* = 0.93, *df* = 7, *p* = 0.38; L2/3, *t* = 2.58, *df* = 8, *p* = 0.033; L5/6, *t* = 2.16, *df* = 4.8, *p* = 0.086; WM, *t* = 2.07, *df* = 8, *p* = 0.072; two-sided Student’s *t*-test or Welch test [L5/6]) and *VIM*+ endothelial cells (L1, *t* = 1.20, *p* = 0.27; L2/3, *t* = 1.47, *p* = 0.18; L5/6, *t* = 0.74, *p* = 0.48; WM, *t* = 0.23, *p* = 0.83; two-sided Student’s *t*-test, *df* = 8 except for 7 for L1) in each cortical layer. **h**, The numbers of *SERPINA3* mRNA molecules in each *SERPINA3*+ astrocyte (L1, 1809 and 2697 cells, *median* = 10.1 and 6.8, *p* < 0.0001; L2/3, 1117 and 1906 cells, *median* = 11.4 and 7.5, *p* < 0.0001; L5/6, 1487 and 2834 cells, *median* = 8.0 and 8.0, *p* = 0.46; WM, 3521 and 6239 cells, *median* = 7.0 and 10.3, *p* < 0.0001, two-tailed Wilcoxon). Asterisk in blue and black indicate a decreasing and increasing trend in SCZ, respectively. **i**, The cellular density (/μm^2^) of *SERPINA3*+ astrocytes (L1, *t* = 0.43, *p* = 0.68; L2/3, *t* = 0.47, *p* = 0.65; L5/6, *t* = 0.81, *p* = 0.44; WM, *t* = 1.20, *p* = 0.26; two-sided Student’s *t*-test, *df* = 8 except for 7 for L1). **j**, The density of *SERPINA3* mRNA molecules (/μm^2^) in each cortical layer (L1, *t* = 0.83, *df* = 7, *p* = 0.44; L2/3, *t* = 0.68, *df* = 8, *p* = 0.51; L5/6, *t* = 0.92, *df* = 8, *p* = 0.38; WM, *t* = 1.35, *df* = 4.4, *p* = 0.24, two-sided Student’s *t*-test or Welch test). Mean ± SEM for bar plots. * *p* < 0.05, ** < 0.01, *** < 0.0001, † < 0.1.

**Extended Fig. 3.**
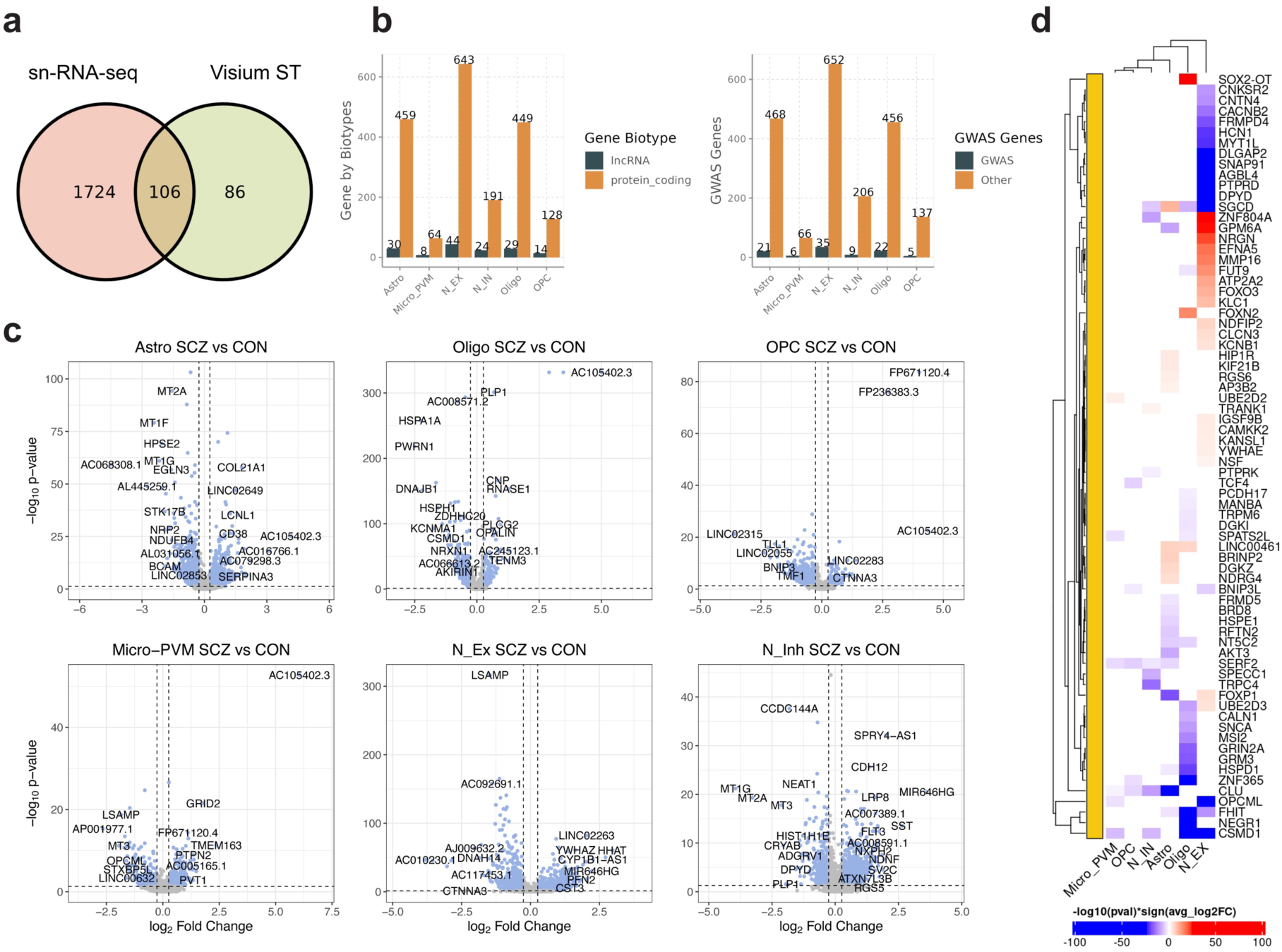
Gene biotypes and GWAS-associated genes dysregulated in SCZ across brain cell types. **a**, Venn diagram showing the number of common and unique genes dysregulated in sn-RNA-seq and Visium ST (*p*_adj_ < 0.05 & FC > 1.2). **b**, Gene biotypes within DGE signatures (*p*_adj_ < 0.05, FC > 1.2) across cell types in SCZ, showing the number of long non-coding and protein-coding genes (left panel) and the number of GWAS-associated genes (right panel). **c**, Volcano plots illustrating the top differentially expressed genes in SCZ compared to CON subjects across cell types, with key upregulated and downregulated genes labeled (*p*_adj_ < 0.05 & FC > 1.2). **d**, Heatmap depicting identified GWAS genes within DGE signatures (*p*_adj_ < 0.05, FC > 1.2) across cell types in SCZ. Color intensity represents the level of significance.

**Extended Data Fig. 4.**
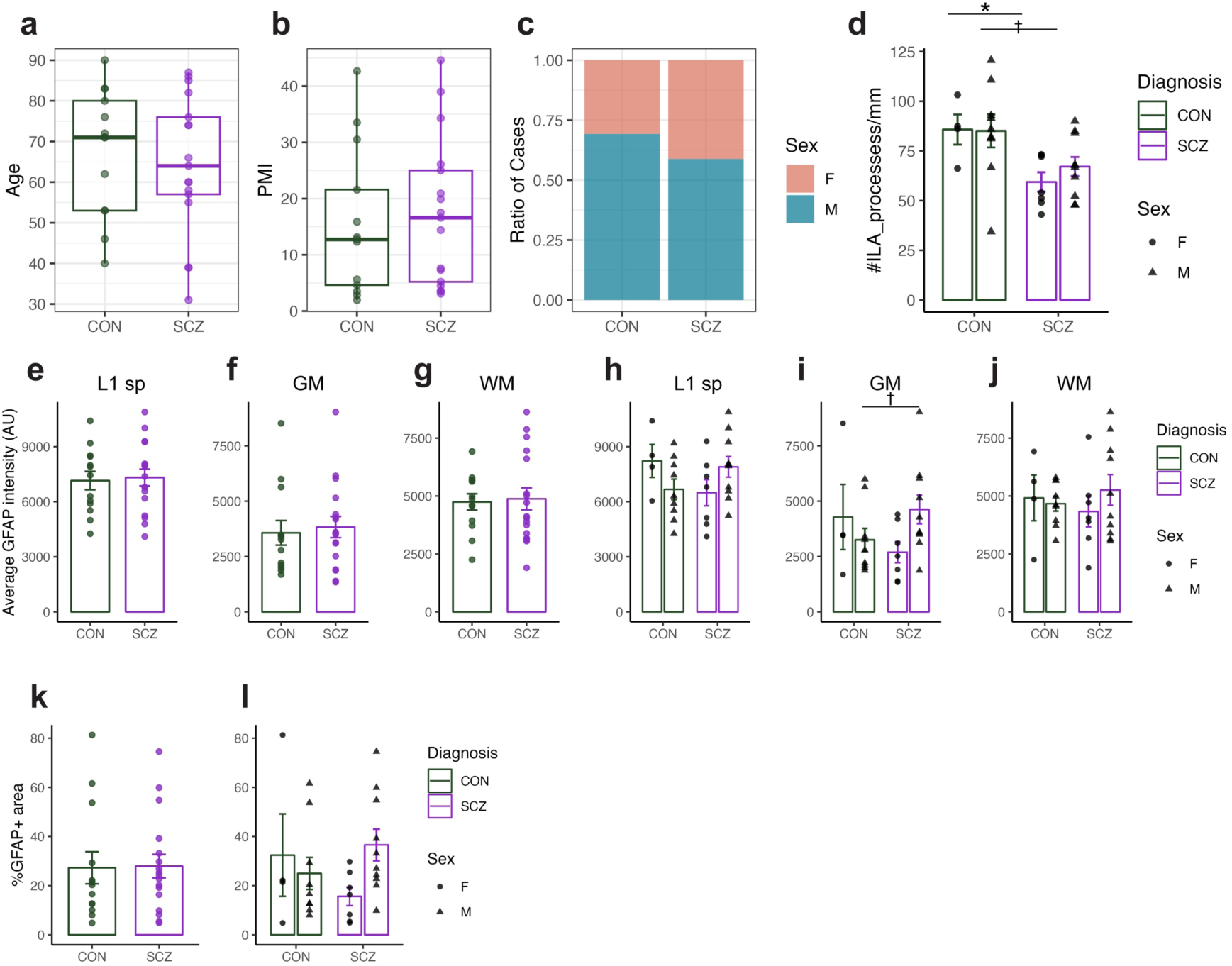
Immunohistochemistry for GFAP in the ACC of SCZ. **a-c**, Demographic characteristics of the 13 CON and 17 SCZ samples used in the immunohistochemical analysis. Age (**a**), postmortem interval (PMI) (**b**), and sex (**c**) do not differ significantly between CON and SCZ (age, *p* = 0.69, two-sided exact Wilcoxon rank-sum test; PMI, *p* = 0.54, two-sided exact Wilcoxon; sex, Fisher’s exact test, *p* = 0.71). **d**, The density of processes of ILAs at 100 μm from the L1/2 boundary, following stratification by sex. The differences between CON and SCZ are significant or marginally significant (9 CON and 10 SCZ male cases, *p* = 0.042; 4 CON and 7 SCZ female cases, *p* = 0.079, two-sided exact Wilcoxon). **e-g**, The average GFAP intensity in the L1 (**e**), GM (**f**), and WM (**g**). No significant differences are observed between CON and SCZ (13 CON and 17 SCZ cases, L1, *p* = 0.80; GM, *p* = 0.51; WM, *p* = 0.90, two-sided exact Wilcoxon). **h-j**, The average GFAP intensity after sex stratification in the L1 (**h**), GM (**i**), and WM (j). The GM shows a marginally significant trend toward increased intensity in SCZ (4 CON vs. 7 SCZ female cases and 9 CON vs. 10 SCZ male cases; L1 female, *p* = 0.23; L1 male, *p* = 0.16; GM female, *p* = 0.41; GM male, *p* = 0.079; WM female, *p* = 0.53; WM male, *p* = 0.78; two-sided exact Wilcoxon). **k-l**, The percentage of area with GFAP immunoreactivity before (**k**) and after (**l**) sex stratification. The GM shows a weak, non-significant trend toward increased intensity in male SCZ (13 CON and 17 SCZ cases of both sexes, *p* = 0.62; 4 CON and 7 SCZ female cases, *p* = 0.65; 9 CON and 10 SCZ male cases, *p* = 0.16; two-sided exact Wilcoxon). * *p* < 0.05, † < 0.1. Mean ± SEM.

**Extended Data Fig. 5.**
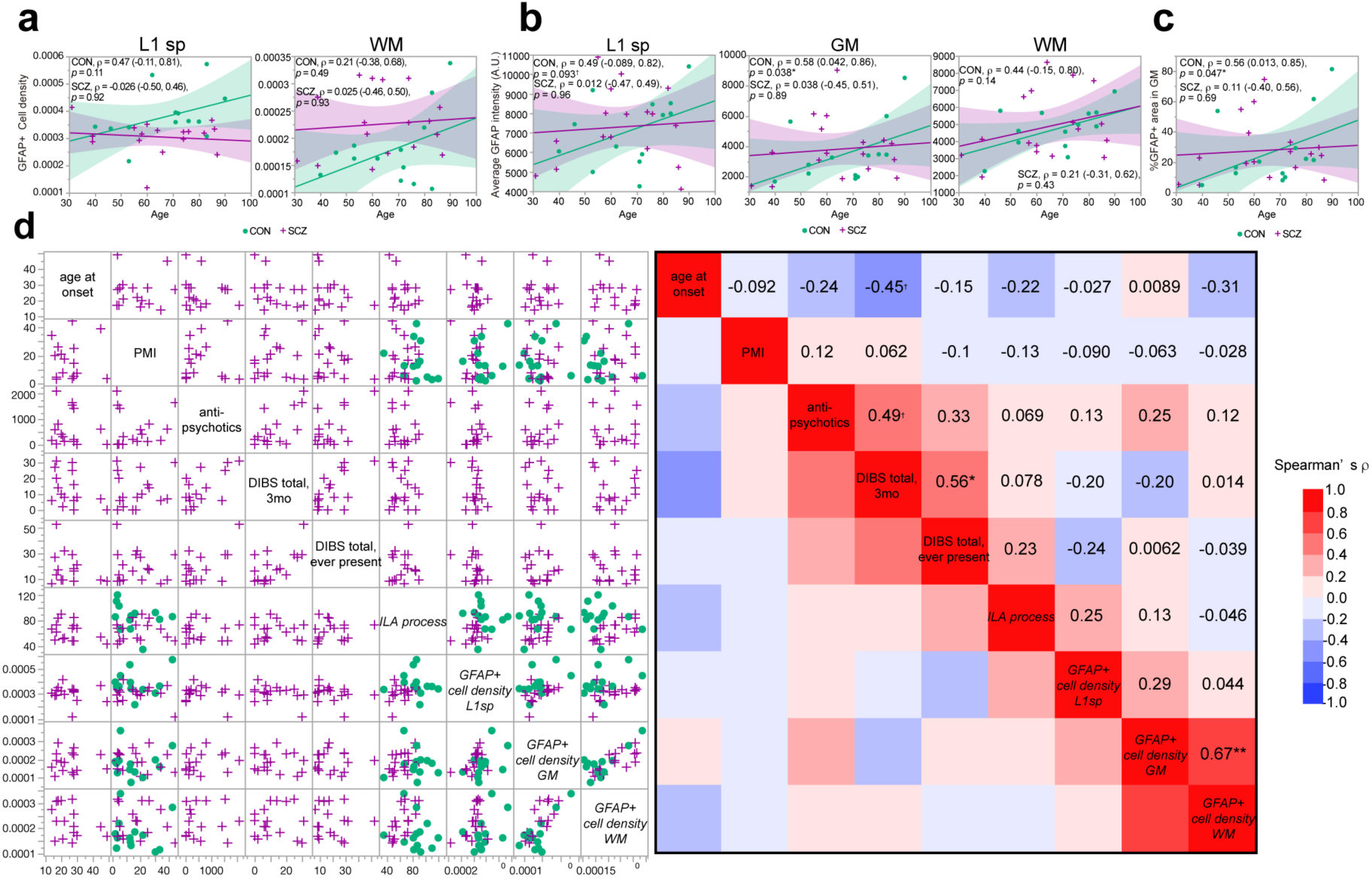
GFAP+ astrocytes in aging and schizophrenia. **a**, Correlation analyses for the L1sp and WM across CON and SCZ groups. Refer to the Fig. 5c, middle panel, for GFAP+ cell density in the GM. The parameters of Spearman correlation [*ρ* (95% confidence interval), *p* value] are shown in the graphs. Linear regression lines with a 95% confidence interval are shown. **b**, Correlation analysis showing the relationship between age and the GFAP fluorescence intensity across CON and SCZ subjects, indicating a positive trend in CON (13 subjects, Spearman) and no significant relationship in SCZ subjects (17 subjects, Spearman). **c**, Correlation analysis showing the relationship between age and the percentage of areas positive for GFAP across CON and SCZ subjects, indicating a positive trend in CON (13 subjects, Spearman) and no significant relationship in SCZ subjects (17 subjects, Spearman). **d**, Associations with representative clinical characteristics and *astrocytic parameters* (italicized). *ρ* is shown in the color map. The age at onset and DIBS total scores in the three months before death (17 subjects, *ρ* = – 0.45 (–0.70, –0.11), *p* = 0.081, Spearman), and the dose of antipsychotics (CP-eq) and DIBS total scores (16 subjects, with CP-eq from one patient missing, *ρ* = 0.49 (0.17, 0.71), *p* = 0.056, Spearman) shows an association with marginal significance. The DIBS total scores in the three months before death are significantly associated with those ever-present (17 subjects, *ρ* = 0.56 (0.26, 0.76), *p* = 0.021, Spearman). The GFAP+ cell density in the GM is associated with that in the WM (30 subjects, *ρ* = 0.67 (0.40, 0.83), *p* < 0.001, Spearman). No significant associations are found between clinical characteristics and *astrocytic parameters*. * *p* < 0.05, ** < 0.001, ***< 0.0001, † < 0.1.

**Extended Data Fig. 6.**
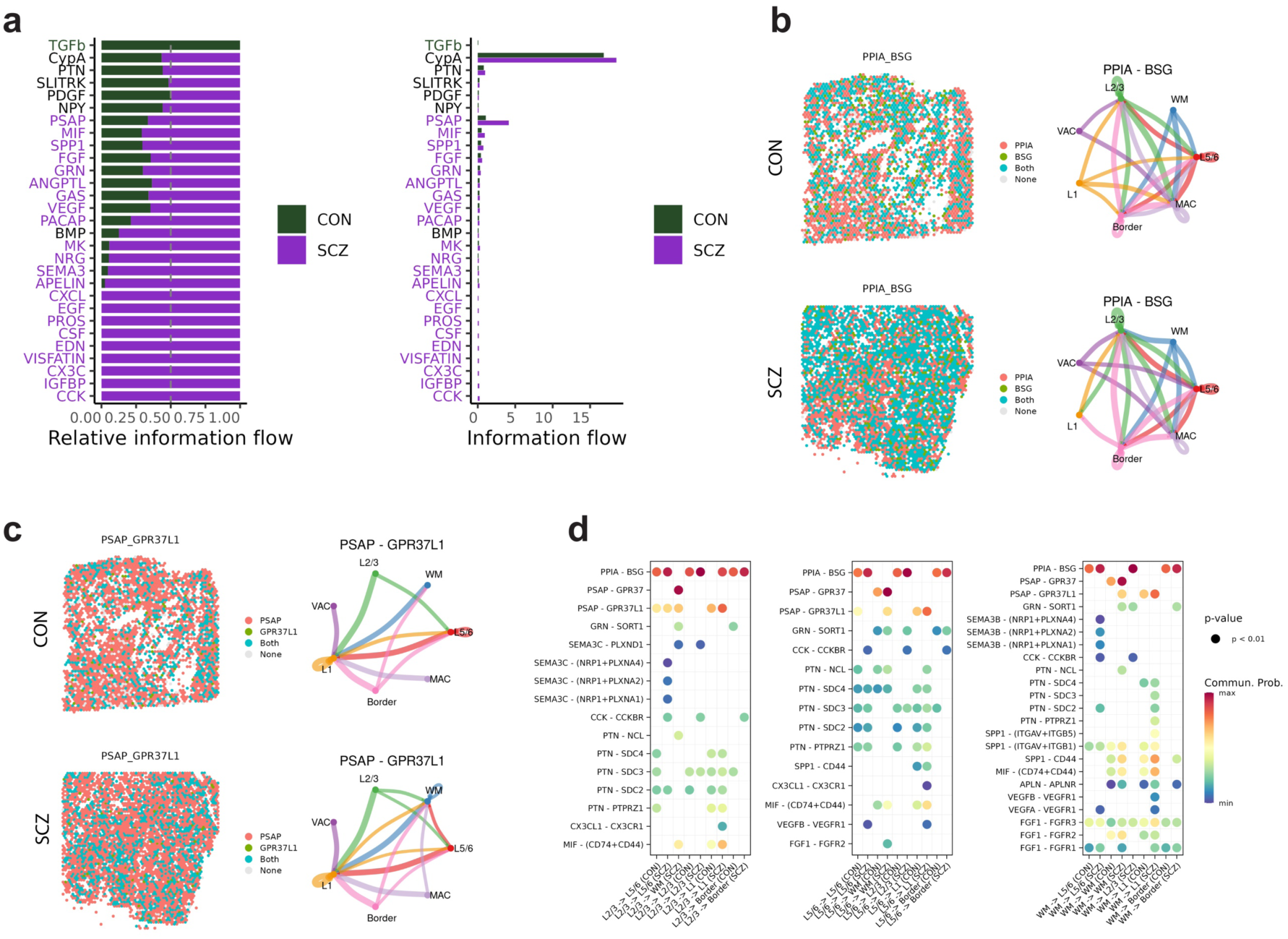
Predominant signaling and L-R interactions in the ACC of CON and SCZ donors. **a**, A comparison of the overall information flow of each signaling pathway in the CON and SCZ groups revealed prominent differences. The top signaling pathways colored purple were enriched in SCZ, while those colored green were enriched in CON. These pathways were ranked based on the differences in the overall information flow within the inferred networks between the two groups. **b**, Spatial distribution of PPIA_BSG, main contributors to the CypA signaling in CON and SCZ, illustrating alterations in signaling localization (left panel). Chord diagrams showing differences in predicted CypA regional communication (right panel). **c**, Spatial distribution of PSAP_GPR37L1, main contributors to the PSAP signaling in CON and SCZ, illustrating alterations in signaling localization (left panel). Chord diagrams showing differences in predicted PSAP regional communication (right panel). **d**, Regional signaling diagrams indicating dysregulated L-R signaling interactions in SCZ, with dot colors denoting statistical significance (*p* < 0.01).

**Extended Data Table 1.** Demographic summary of brain samples analyzed in this study.

| Diagnosis | Sex | Age at death (y/o) | Age at onset (y/o) | PMI (h) | RIN | pH | CP-eq (mg/d) | State | sn-RNA-seq | Visium | RNA-FISH | IHC | Neuropathologist evaluation |
| --- | --- | --- | --- | --- | --- | --- | --- | --- | --- | --- | --- | --- | --- |
| SCZ | F | 87 | 39 | 3.5 | 9.6 | 6.7 | 0 | Fresh Frozen & FFPE | YES | NO | YES | YES | YES |
| SCZ | F | 76 | 29 | 25 | 5.7 | 6.1 | 114 | Fresh Frozen & FFPE | YES | NO | YES* | YES | NO |
| SCZ | M | 66 | 31 | 7.3 | 6.7 | 6.2 | 0 | Fresh Frozen & FFPE | YES | YES | YES | YES | YES |
| SCZ | M | 74 | 22 | 17.5 | 8.9 | 6.6 | 120 | Fresh Frozen & FFPE | YES | YES | YES | YES | YES |
| SCZ | F | 82 | -** | 16.6 | 8.5 | 6 | - | Fresh Frozen & FFPE | YES | YES | YES | YES | NO |
| SCZ | M | 74 | 20 | 20.9 | 9.3 | 6.1 | 400 | Fresh Frozen & FFPE | YES | YES | YES | YES | NO |
| CON | F | 83 | N/A | 42.7 | 8 | 6.3 | 0 | Fresh Frozen & FFPE | YES | YES | YES | YES | NO |
| CON | M | 76 | N/A | 33.5 | 8.9 | 6.4 | 0 | Fresh Frozen & FFPE | YES | YES | YES | YES | NO |
| CON | M | 53 | N/A | 15.9 | 6.1 | 6.3 | 0 | Fresh Frozen & FFPE | YES | YES | YES | YES | NO |
| CON | F | 74 | N/A | 31.7 | 5.8 | 5.9 | 0 | Fresh Frozen & FFPE | YES | NO | YES | YES*** | NO |
| CON | F | 90 | N/A | 5.6 | 6 | - | 0 | Fresh Frozen & FFPE | YES**** | NO | YES**** | YES | NO |
| CON | M | 83 | N/A | 30.5 | 6 | - | 0 | Fresh Frozen & FFPE | YES | YES | YES | YES | NO |
| SCZ | F | 31 | 21 | 7.6 | - | - | 305 | FFPE | NO | NO | NO | YES | NO |
| SCZ | M | 64 | 17 | 4.5 | - | - | 15 | FFPE | NO | NO | NO | YES | NO |
| SCZ | M | 60 | 27 | 44.6 | - | - | 1650 | FFPE | NO | NO | NO | YES | NO |
| SCZ | F | 39 | 16 | 19.9 | - | - | 580 | FFPE | NO | NO | NO | YES | YES |
| SCZ | F | 57 | 17 | 3.1 | - | - | 2109 | FFPE | NO | NO | NO | YES | NO |
| SCZ | M | 39 | 14 | 34.3 | - | - | 155 | FFPE | NO | NO | NO | YES | NO |
| SCZ | M | 58 | 28 | 5.2 | - | - | 1427 | FFPE | NO | NO | NO | YES | NO |
| SCZ | F | 85 | 45 | 39 | - | - | 606 | FFPE | NO | NO | NO | YES | NO |
| SCZ | M | 55 | 27 | 3.6 | - | - | 1550 | FFPE | NO | NO | NO | YES | NO |
| SCZ | M | 60 | 18 | 26.1 | - | - | 800 | FFPE | NO | NO | NO | YES | NO |
| SCZ | M | 86 | 28 | 14.4 | - | - | 200 | FFPE | NO | NO | NO | YES | NO |
| CON | M | 71 | N/A | 2.8 | - | - | 0 | FFPE | NO | NO | NO | YES | YES |
| CON | M | 71 | N/A | 2.0 | - | - | 0 | FFPE | NO | NO | NO | YES | YES |
| CON | M | 53 | N/A | 3.5 | - | - | 0 | FFPE | NO | NO | NO | YES | NO |
| CON | F | 80 | N/A | 4.6 | - | - | 0 | FFPE | NO | NO | NO | YES | YES |
| CON | M | 62 | N/A | 12.3 | - | - | 0 | FFPE | NO | NO | NO | YES | YES |
| CON | M | 72 | N/A | 12.7 | - | - | 0 | FFPE | NO | NO | NO | YES | YES |
| CON | F | 40 | N/A | 13.1 | - | - | 0 | FFPE | NO | NO | NO | YES | YES |
| CON | M | 46 | N/A | 21.6 | - | - | 0 | FFPE | NO | NO | NO | YES | NO |

SCZ, schizophrenia; CON, control; F, female; M, male; PMI, postmortem interval; RIN, RNA integrity number; CP-eq, chlorpromazine equivalent doses of antipsychotics 3 month before death
\*excluded from the study due to poor staining of markers.
\*\*developed symptoms in her teenage, but precise age at onset unknown.
\*\*\*excluded from the study because it did not contain the full thickness of the cortex.
\*\*\*\*excluded from the analysis because the total cell density was exceptionally high in histological analysis.

